# Strobemers: an alternative to k-mers for sequence comparison

**DOI:** 10.1101/2021.01.28.428549

**Authors:** Kristoffer Sahlin

**Affiliations:** Department of Mathematics, Science for Life Laboratory, Stockholm University, 106 91, Stockholm, Sweden

**Keywords:** k-mers, minimizers, sequence matching, data structures

## Abstract

K-mer-based methods are widely used in bioinformatics for various types of sequence comparison. However, a single mutation will mutate *k* consecutive k-mers and makes most k-mer based applications for sequence comparison sensitive to variable mutation rates. Many techniques have been studied to overcome this sensitivity, *e.g*., spaced k-mers and k-mer permutation techniques, but these techniques do not handle indels well. For indels, pairs or groups of small k-mers are commonly used, but these methods first produce k-mer matches, and only in a second step, a pairing or grouping of k-mers is performed. Such techniques produce many redundant k-mer matches due to the size of *k*.

Here, we propose *strobemers* as an alternative to k-mers for sequence comparison. Intuitively, strobemers consist of linked minimizers. We use simulated data to show that strobemers provide more evenly distributed sequence matches and are less sensitive to different mutation rates than k-mers and spaced k-mers. Strobemers also produce a higher match coverage across sequences. We further implement a proof-of-concept sequence matching tool StrobeMap, and use synthetic and biological Oxford Nanopore sequencing data to show the utility of using strobemers for sequence comparison in different contexts such as sequence clustering and alignment scenarios. A reference implementation of our tool StrobeMap together with code for analyses is available at https://github.com/ksahlin/strobemers.

## Introduction

The dramatic increase in sequencing data generated over the past two decades has prompted a significant focus on developing computational methods for sequence comparison. A popular sequence comparison paradigm is k-mer based analysis, where k-mers are substrings of length *k* of, *e.g*., genomic, transcriptomic, or protein sequences. K-mer-based methods have been applied for sequence comparison for error correction (1), genome assembly (2, 3), metagenomic (4) and chromosome (5) sequence classification, sequence clustering (6), database search (7, 8), structural variation detection (9–11), transcriptome analysis (12, 13), DNA barcoding of species (14), estimation of genome size (15), identification of biomarkers (16), and many other applications. Because of the widespread use of k-mers, many data structures and techniques for efficiently storing and querying k-mers have been proposed (see (17) for a review).

While k-mers has proven to be practical in several sequence comparison problems, they are sensitive to mutations. A mutation will mutate *k* consecutive k-mers across a string. As the mutation rate increases, the number of matching k-mers between two sequences quickly reduces. In (18), the distribution of mutated k-mers was studied in detail. The authors provided closed-form expressions for the mean and variance estimates on the number of mutated k-mers under a random mutation model. While the number of k-mer matches between sequences is of interest, it is often more informative to know how they are distributed across the matching region. K-mer matches, because of their consecutive nature, cluster tightly in shared sequence regions, while matches may be absent in regions with higher mutation rates. Spaced k-mers (or spaced seeds) have been studied in several sequence comparison contexts to overcome the k-mers’ sensitivity to mutations (19–21). The advantage of spaced k-mers is that matches of spaced k-mers at different positions are less correlated with each other than k-mer matches. In fact, k-mers are in some conditions the worst seed pattern for the problem of similarity search (22). Another innovative idea has been to permute the letters in a string before comparison (23, 24). The main idea is to permute the letters in regions of fixed size in a string using several different permutations. When comparing two strings in the regions under these permutations, at least one permutations will, with statistical certainty, have pushed any substitution(s) towards the end of the region. This allows for a constant-time query of the prefix of the region in the permuted strings. With more permutations, it is more likely to find an exact prefix match. However, both spaced k-mers and permutation techniques are only practical for substitutions. An insertion or deletion (*indel*) will shift the sequence and, similarly to k-mers, result in a long stretch of dissimilar k-mers. For certain applications such as genome assembly, selecting several sizes of *k* for inference has also been shown to help sequence comparison (25), but it significantly increases runtime and complexity of analysis. There are also methods to collapse repetitive regions before k-mer based comparison (26), which reduces the processing time of repetitive hits. However, such techniques are usually employed for reference-based analysis and do not apply to general sequence comparison problems.

As third-generation sequencing techniques appeared with sequencing errors mostly consisting of insertions and deletions, many of the previously developed sequence comparison techniques for short-read data became unsuitable. For the third generation sequencing, MinHash (27) and minimizers (28, 29) proved to be useful data structures for such sequence comparison as minimizers can be preserved in a window affected by an indel. In addition, they also reduce the size of the index by subsampling the data. This has made MinHash and minimizers a popular technique for subsampling k-mers employed for sequence comparison in a range of applications such as metagenome distance estimation (30) and alignment (31, 32), clustering (33) error-correction (34), and assembly (35) of long-read sequencing data. Because of the widespread practical use of minimizers, several methods have been proposed for sampling them with as low density as possible (36) and more evenly (37, 38).

Due to the error rates of long-read sequencing, minimizers are often chosen much shorter (about 13-15nt) than what is considered to produce mostly unique k-mers in, e.g., the human genome (around *k* > 20nt). With this length of minimizers, they produce many candidate sequence matches. Therefore, it would be useful to combine the robustness of minimizers to indels and mutation errors with larger k-mer sizes that would offer more unique matches. One approach is to use a small k-mer size and identify pairs (39) or groups (40) of them clustered tightly together, and it has been studied how to design the sampling distribution of seeds to optimize alignment sensitivity (41, 42). Multi-seed methods are robust to any mutation type and have shown to, e.g., improve overlap detection between long reads (43). However, they still match single k-mers individually and group them based on statistics after individual k-mer hits have been found. To remove the redundancy in matches, we suggest that it is beneficial to couple the k-mers in the initial matching step and perform a single constant-time lookup of coupled k-mers. Coupled k-mers have been explored in, e.g., (34, 44) where *paired minimizers* are generated and stored as a single hash. A paired minimizer-match signals that the region is similar between sequences. Due to the gap between the minimizers, such a structure is not as sensitive to indels or substitutions as k-mers. Paired minimizers were shown to be useful for both genome assembly (44) and error correction of long cDNA reads (34) where the reads are similar only in some regions due to alternative splicing. However, in both (44) and (34), minimizers are produced independently and paired up after the minimizer generation. Here, we show that we can substantially improve on paired minimizers for sequence matching by producing minimizers chosen jointly depending on previous windows. We also generalize paired minimizers to link more than two together.

We propose a novel method to extract subsequences from a string, and we call those subsequences *strobemers*. The name is inspired by strobe sequencing technology (an early Pacific Biosciences sequencing protocol), which would produce multiple subreads from a single contiguous fragment of DNA where the subreads are separated by ‘dark’ nucleotides whose identity is unknown, illustrated in (45). Strobemers introduced here are, however, produced computationally. Intuitively, strobemers are groups of linked short k-mers (called *strobes*) from adjacent windows. The strobes can be chosen as minimizers either independently within windows, which we call *minstrobes*, or dependent on previous strobes under two different functions, called *randstrobes*, and *hybridstrobes*.

We show that strobemers (particularly randstrobes and hybridstrobes) have a significant benefit over k-mers and spaced k-mers. Strobemers improve sequence matching by providing more evenly distributed matches than k-mers, are less sensitive to different mutation rates and give a higher total coverage of matches across strings. We also show on human chromosomes that strobemers can offer higher uniqueness, hence confidence in a match, than k-mers due to the spacing of the strobes. We use synthetic and biological Oxford Nanopore sequencing reads to show that strobemers produce more contiguous matches and better coverage when mapping reads to themselves or to a reference sequence. Strobemers are easy to both construct and query, making them a compelling alternative to k-mers and spaced k-mers for sequence comparison. We show that while randstrobes have both a higher theoretical and practical runtime over generating k-mers, minstrobes and hybridstrobes have the same practical runtime as computing minimizers. Furthermore, strobemers can, similarly to k-mers and spaced k-mers, be subsampled using a thinning protocol such as MinHash, minimizers, or syncmers (38).

## Methods

### Definitions

We refer to a *subsequence* of a string as a set of ordered letters that can be derived from a string by deleting some or no letters without changing the order of the remaining letters. A substring is a subsequence where all the letters are consecutive. We use *i* to index the position in a string *s* and let *s*[*i* : *i* + *k*) denote a *k-mer* substring at position *i* in *s* covering the *k* positions *i*,…,*i* + *k* – 1 in *s*. We will consider 1-indexed strings. If *s*[*i*: *i* + *k*) is identical to a k-mer *t*[*i*′: *i*′ + *k*) in string *t*, we say that the k-mers *match*, and that the match occurs at position *i* in *s* (and *i* in *t*). Similarly, let *f* (*i,k,s*) be any function to extract a subsequence of length *k* with first letter at position *i* from *s*. If *f* (*i,k,s*) is identical to *f* (*i*′,*k,t*), we say that the subsequences match, and that the match occurs at position *i* in *s* (and *i* in *t*). For example, for k-mers we have *f* (*i, k,s*) = *s*[*i*: *i* + *k*). We let |·| denote the length of strings.

We use *h* to denote a hash function 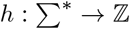 mapping strings to random numbers. Given two integers *w* > 0 and *k* > 0, the minimizer at position *i* in *s* is the substring of *s* of length *k* starting in the interval of [*i, i* + *w*) that minimizes *h*.

### Aim

We will introduce strobemers by describing the problem they aim to solve. Consider two strings *s* and *t* that are identical up to *m* mutations. We desire a function *f* to produce a set of subsequences from *s* and *t* that have two characteristics: (1) as few possible placements of the *m* mutations result in no matches between *s* and *t*, and (2) the sub-sequences of length *k* should be as unique as k-mers on *s* and *t*. Characteristic 1 and 2 relate to match sensitivity and precision, and we will discuss this in an example below. For practical purposes, we also require that at most one subsequence is produced per position to mimic how k-mers are derived in a string (and limit the amount of data we store for each string). Certainly, we could produce all possible subsequences at each position to minimize criteria 1, but this is not feasible. A similar objective to characteristic (1) was studied for multi-seed design (41), where the authors want to find a set of seeds so that at least one seed matches a gapless alignment between two sequences.

### A motivational example

Consider two strings of 100 nucleotides with *m* = 3 mutations between them. This could occur, e.g., in splice alignment to an exon, or in sequence clustering. If we use a k-mer of size 30 to find matches, and the two strings differ at positions 25, 50, and 75, there will be no matching k-mers. Similarly, this holds for mutations at positions 20, 48, and 73 and several other combinations. As described, we want as few possible placements of errors leading to the region being unmatched.

Using spaced k-mers (19) or permutations of the string (24) would help if the mutations were substitutions. We could consider lowering *k*, but this would generate more matches to other strings as well. To achieve the same uniqueness as longer *k*, we could consider coupled k-mers (39) of say 15nt per pair, with some gap in between them. Note that the k-mers would need to be coupled before searching for matches to avoid many matches to other sequences. Furthermore, if the coupled k-mers have a fixed distance from each other, we have just created a specific type of spaced k-mers, which are only robust to substitutions. We, therefore, could consider coupled minimizers (34, 44) to select a random gap size for us, but in a deterministic manner.

This brings us to the strobemers. In the scenario above, we could pick a k-mer of size 15 at a position we want to sample and couple it with a minimizer of length 15 derived from a window downstream from the k-mer. Together, they have sequence length 30 and are therefore robust to false matches. They can also match across the mutations, where the mutations could be both substitutions and indels. If we increase the mutation density on our string, eventually, our two k-mers of length 15nt will also fail to produce any matches. Therefore, we could consider triplets of a k-mer and two minimizers of length, e.g., 10nt. Finally, we can further reduce the sampled minimizers’ dependency, and therefore the matches, using other hashing protocols (as we will investigate here).

### Strobemers

Consider a string *s*. A strobemer of order *n* in *s* is a subsequence of *s* composed of a set of ordered substrings *m*_1_,…*m_n_* on *s* of equal length ℓ, that we call *strobes*. If the first strobe *m*_1_ starts at position *i*, the second strobe *m*_2_ will be chosen from a window [*i* + *w_min_*: *i* + *w_max_*] with *w_min_* < *w_max_* on *s*, *m*_3_ from [*i* + *w_min_* + *w_max_* : *i* + 2*w_max_*], and *m_n_* from [*i* + *w_min_* + (*n* – 2)*w_max_* : *i* + (*n* – *1*)*w_max_*]. We will from now on parametrize a strobemer as (*n,ℓ,w_min_,w_max_*) denoting the order, the length of the strobe, and the minimum and maximum window offset to the previous window, respectively.

We select *m*_2_,…*m_n_* based on some hash function. We will consider three different hash functions for producing them, which give significantly different results. First we denote as *minstrobe*, a strobemer where strobes *m*_2_,…,*m_n_* are independently selected as minimizers in their respective windows under a hash function *h* (Figure 1A).

**Fig. 1.**
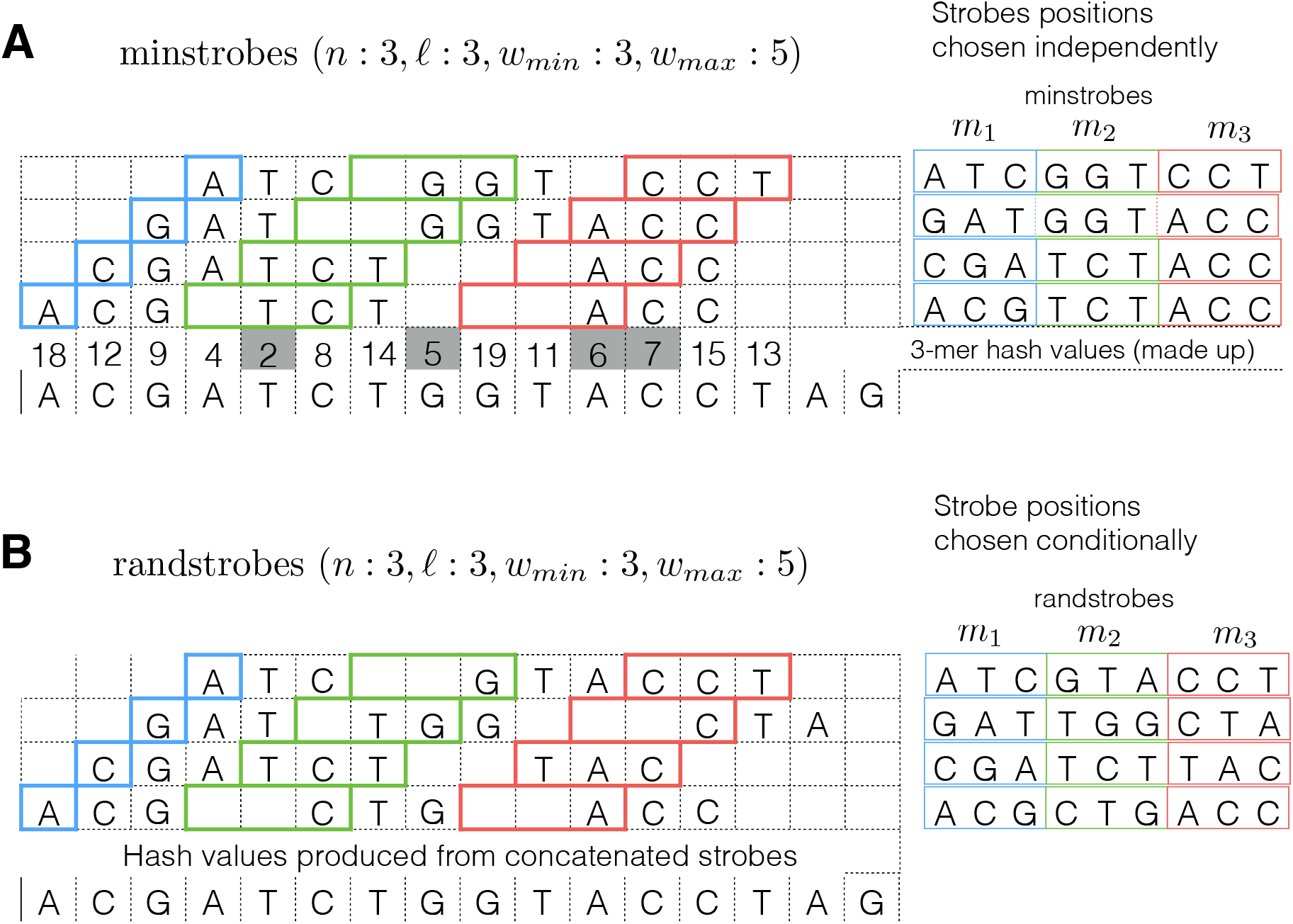
An illustration of four minstrobes (A) and randstrobes (B) with (*n* = 3, ℓ = 3, *w_min_* = 3, *w_max_* = 5) generated from a DNA string of 16 letters. With parameters *n* = 3 and ℓ = 3, the strobemers will consist of three strobes (substrings) each of length 3. The position of the first strobe *m*_1_ in each of the four strobemers is highlighted in blue. The rest of the strobemers are chosen from a window of *w_max_* – *w_min_* + 1 = 3 positions based on a minimizer protocol of minstrobes (A) or randstrobes (B). The possible start positions of strobes *m*_2_ and *m*_3_ are highlighted in green and red, respectively. In the minstrobe protocol (A), the 3-mer minimizer hash values (under a made up hash function in the figure) are showed above the DNA string and come from computing *h*(*m*) for each 3-mer strobe *m*. The position of the hash value corresponds to the first position of the 3-mer strobe. The minimizer values in all relevant strobe windows of length 3 in the figure are indicated by grey squares. In the minstrobe protocol, strobes *m*_2_ and *m*_3_ are selected independently based on the minimizer value in each strobemer window. This gives a high similarity between nearby strobemers (sharing minimizers). The four minstrobes produced are shown to the right in A. In the randstrobe protocol (B), strobes *m*_2_ and *m*_3_ are selected dependent on the previous strobes, *i.e*., *h*(*m*|*m*_1_,…,*m*_*i*–1_). The function producing the conditional dependence is irrelevant for the purpose of illustration. Here we use string concatenation of previous strobes to produce the dependence, but any other function producing conditional dependence will suffice. Because of the conditional dependence in the hash function, randstrobes are more randomly (but deterministically) distributed across the sequence.

Second, we denote as *randstrobe*, a strobemer where strobe *m_j_* is selected as minimizer dependent on the previous *m*_1_,…*m*_*j*−1_ strobes (Figure 1B). Any asymmetric hash function (*i.e*, *h*(*s,t*) ≠ *h*(*t,s*)) with conditional dependence on previous strobes suffice for our purposes in this study. Here, we chose the hash function *h*(*m*′|*m*_1_,…,*m*_*j*−1_) = *h*(*m*_1_ ⊕ … ⊕ *m*_*j*–1_ ⊕ *k*′), where φ denotes string concatenation, where *h* concatenates the previous selected strobes *m*_1_,…,*m*_*j*−1_. Thus, the k-mers in the randstrobe are produced iteratively from *i = 1,..,n* and yields a more randomly distributed set of strobes.

Third, we will consider a hybrid between minstrobes and randstrobes that uses both independent minimizers and a conditional hash function that we call *hybridstrobes*. Consider partitioning the sampling window for each strobe into *x* disjoint segments of length *w_x_* = ⌊(*w_max_* – *w_min_*)/*x*⌋. That is the sampling window for *m*_2_ is partitioned into [*i* + *w_min_* : *i* + *w_min_* + *w_x_*) [*i* + *w_min_* + *w_x_* : *i* + *w_min_* + 2*w_x_*),… [*i* + *w_min_* + (*x* – 1)*w_x_* : *i* + *w_max_*), and similarly for the sampling windows of the other strobes. We select a strobe *m_j_* as the minimizer in the *r*th window segment of length *w_x_* dependent on the remainder *r* of the previous strobe modulo *x*, i.e., *r* = *h*(*m*_*j*–*i*_) mod *x*. While this selection is not as randomly distributed as randstrobes, the variability of *x* possible pairings provides more randomly distributed matches than minstrobes. Here we will use *x* = 3.

There are two important aspects to consider for the three protocols. Firstly, for two strobes with nearby starting positions, strobes *m*_2_,..,*m_n_* will most frequently be the same under the minstrobe generation due to independent minimizers (see Fig. 1), and most frequently differ in a randstrobe. This means that under the same parameters in the protocols, the randstrobes will (in all likelihood) contain more uniquely sampled positions and, hence, more unique randstrobes, while minstrobes more frequently share minimizers. Hybridstrobes places somewhere in between minstrobes and randstrobes depending on the size of *x*.

Secondly, generating minstrobes and hybridstrobes are in practice almost as fast as producing minimizers while generating randstrobes, under the function we consider here, is not. We elaborate on this in the section on time complexity. Finally, we note that minstrobes of order 2 are similar to but formally different from paired minimizers (34, 44). Both minstrobes of order 2 and paired minimizers consist of two substrings. However, paired minimizers are two minimizers that are coupled together under some distance constraint on a sequence. In the minstrobe protocol, the first strobe is not necessarily a minimizer. However, strobemers can be subsampled with a thinning protocol. In this case, a strobemer with *n* = 2 can be considered as a specific method to select paired minimizers.

### Construction of strobemers

We aim to produce a strobemers of a string *s* in a similar manner to how k-mers are produced, *i.e*., one strobemer per position *i* ∈ [1, |*s*| – *k* + 1]. This would mean that we extract the same amount of k-mers and strobemers from a string *s*, and consequently, for equal length *k*, the same amount of raw data. Note however, that the number of unique k-mers and strobemers may differ. We construct strobemers as follows. The total possible span of a strobemer of order *n* is *W* = (*n* – 1)*w_max_* + ℓ, and the total subsequence length as 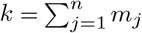 (no strobe is overlapping). Let us consider extraction of a strobemer at position *i* in *s*. If *W* ≤ |*s*| – *i* we use the predefined windows [*w_min_, w_max_*) and compute the strobemers under the respective strobemer protocols as described above. If *W* > |*s*| – *i*, we narrow the window sizes until *m*_1_ to *m_n_* are all adjacent to each other producing a substring (k-mer) of length *k*. Under this construction, the same amount of k-mers and strobemers will be extracted from a string. Any way to narrow the windows at the end of the sequence can be considered. Here, we choose to shorten each window [*w_min_,w_max_*] to 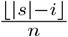. Furthermore, while the protocol to extract strobemers allows overlapping strobes, here we will only consider *w_min_* ≥ ℓ giving non-overlapping strobes. Pseudocode to construct strobemers are given in appendix A.

### Time complexity

If we ignore the time complexity of the hash function, the time complexity of generating minimizers is *O*(|*s*|*w*) for a window size *w*. However, as (31) noted, computing minimizers is in practise close to *O*(|*s*|) if we use a queue to cache previous minimizer values in the window. The expensive step is when a previous minimizer is discarded from the queue and a new minimizer needs to be computed for the window.

Similar to computing minimizers, strobemers have the worst-case time complexity of *O*(|*s*|*n*(*w_max_* – *w_min_*)). However, the independence of hash values in the minstrobe and hybridstrobe protocols makes them close to *O*(|*s*|) in practice by using separate queues for each strobe sampling window in the same manner as computing minimizers independently. The randstrobe protocol does not have this independence under the hashing scheme we consider in this study, which means that all hash values have to be recomputed at each position. This means its practical time complexity is therefore *O*(|*s*|*n*(*w_max_* – *w_min_*)).

### Implementation

The pseudocode to construct strobemers (appendix A) are provided for the simplicity in expression, they are not efficient implementations. We want to avoid string concatenation. We also want to avoid repeated computation of minimizers for minstrobes and hybridstrobes where minimizer values are computed independently.

For minstrobes and hybridstrobes, we first precompute all the hash values in a string to work with addition of hash values and not string concatenations. For a minstrobe or hybridstrobe of order 2 consisting of strobes *m*_1_ and *m*_2_, the strobemer hash value that is stored will be *h*(*m*_1_) – *h*(*m*_2_). We store the hash values to represent the strings and not the strings explicitly. Similarly, for a minstrobe or hybridstrobe of order 3 consisting of strobes *m*_1_, *m*_2_, and *m*_3_, we store *h*(*m*_1_) – *h*(*m*_2_) + 2*h*(*m*_3_). An asymmetric function should be used so that a permutation of the strobemers do not produce the same hash values. As described in (31) We also keep a queue datastructure for each strobe *m_j_*, *j* ≥ 2 and the current minimum hash value in these windows so that we only need to recompute the minimum hash values in the window whenever we discard the current minimum in the queue.

For randstrobes, we similarly to the other protocols, precompute all the hash values in a string and store only the hash values representing randstrobes. Therefore, instead of taking the minimum over string concatenations in a window as described in the pseudocode for randstrobes (appendix A), we select the strobe *m_j_* in a window that minimizes the function *h*(*m*) – *h*(*m_j_*) mod *q*, where *q* is a prime (we choose 997). Similarly to minstrobes, the final hash value to represent the randstrobe is *h*(*m*_1_) – *h*(*m*_2_) for *n* = 2 and *h*(*m*_1_) – *h*(*m*_2_) + 2*h*(*m*_3_) for *n* = 3.

## Results

### Overview

We will first investigate sequence matching performance of strobemers (order 2 and 3) compared to k-mers and spaced k-mers using simulated data. We consider both how effective the different protocols are at finding matches under different error rates (related to sensitivity) and how unique the matches are that they produce (related to specificity).

We then implement a tool StrobeMap, and use synthetic and biological data to demonstrate the utility of strobemers in various applications. We map ONT cDNA reads with 7% median error rate from (34) both to themselves and to reference sequences. We also map genomic ONT *E. coli* reads with 17% median error rate both to themselves and to an *E. coli* genome, as well as two *E. coli* genomes to themselves. In the experiments we compare the contiguity and coverage of the matches produced by k-mers and strobemers.

### Experiment design

The size of the extracted subsequence length *k* of any protocol is central when comparing the efficacy of finding matches and their uniqueness. Therefore, we are interested in comparing sizes of subsequences that are similar between the protocols. Specifically, if the size of the k-mer is 30, we want to compare the k-mers to strobemers parameterized, e.g., by (2, 15, ·,·) and (3, 10, ·,·) as all the extracted subsequences have a length of 30 on the strings. The spaced k-mers consists of a window of size *L* with *k fixed* positions and a set of *L* – *k* wildcard (or “don’t care”) positions. This is commonly represented as a binary string where 1’s are sampled and 0’s are wildcard positions. For example, in the string *AGGTCA* with *L* = 6, the spaced k-mer 101011 is *AGCA*. In our evaluations, we choose two densities of fixed positions for the spaced k-mers. First, we denote as *spaced-dense* a strategy where 2/3 of the positions are fixed, and *spaced-sparse* where 1/3 of the positions are fixed. The spaced-dense and the spaced-sparse frequency of fixed positions roughly correspond to the densities used in (20) and (46), respectively. To keep *k* fixed, *L* = 1.5*k* in the spaced-dense protocol and *L* = 3*k* in the spaced-sparse protocol. The windows’ first and last positions are always fixed (as in (20, 46)) to assure the length of the spaced k-mer. The remaining fixed positions are randomly chosen. In, e.g., (46), the sampled positions are handpicked. While handpicking positions may be more suitable for optimizing lower correlation between matches, this study focuses on designing a protocol robust to indels. We will observe that spaced k-mers do not work well for mutations other than substitutions.

### Evaluation metrics

If a k-mer or spaced k-mer extracted from position *i* in *s* and *i*′ in *t* produce the same hash value, we say that a *match* between two sequences *s*_1_ and *s*_2_ occur at position *i* and *i*′ in the two strings respectively. For a k-mer, we say that the match produces a *sequence coverage* over positions [*i,i* + *k*]. For a spaced k-mer, we say that the match produces a *sequence coverage* over the *k* fixed (sampled) positions. Furthermore, for a k-mer we say that the match has a *match coverage* of length *k* (*i.e*., positions [*i, i* + *k*]), and of length *L* in case of the spaced k-mer (*i.e*., the span of the fixed positions). If a strobemer extracted from position *i* in *s* and *i* in *t* produce the same hash value, we say that a match between two sequences *s*_1_ and *s*_2_ occur at position *i* and *i*′ as well as at the start positions of the additional strobes *m*_2_,…,*m_n_* in the two strings respectively. We say that the match produces a *sequence coverage* over all the positions covered by the strobes in the match. Furthermore, we say that the match has a *match coverage* spanning the first nucleotide in the first strobemer to the last nucleotide in the last strobemer. The total sequence coverage and match coverage of a string *s* is calculated as the union of all positions covered under the definitions of sequence coverage and match coverage, respectively. We adopt similar terminology as in (18) and denote a maximal interval of consecutive positions without matches as an *island*.

To evaluate the sequence matching ability, we compare under different error rates (i) the fraction of matches, (ii) the sequence coverage, (iii) the match coverage, and (iv) the distribution of islands. We need to make two clarifications on these evaluation metrics. First, our experiments on simulated data are designed with parameters so that the event of observing a false match (*e.g*., repetitive k-mer) under any protocol has a negligible probability. This means that our simulated experiments only measure the raw ability to identify correct matches.

Second, as for the distribution of islands, we are interested in measuring the sizes of islands and their size distribution. We calculate the island *E-size* (47), a commonly used metric in genome assembly that we will adapt for our purposes. For a string *s* and a set of islands lengths *X* on *s* we calculate the island E-size *E* as follows.

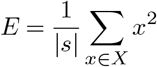

*E* measures the *expected island size*, and intuitively, we can think of *E* as follows. We pick a position at random across *s* and observe the island size spanning that position. We may pick positions that are covered by matches (i.e., island size 0), but if we keep picking positions at random over *s* and store our observations on the island lengths, we will end up with *E* according to the *law of large numbers*. We will also show the entire island distribution.

### Strobemer vs k-mer matching

We compare how effective the different protocols are at producing matches for different error rates. We start with a controlled scenario, where mutations are distributed with a fixed distance. In our second experiment, we use a random mutation distribution. We perform the fixed-distance mutation experiment to illustrate the advantage of strobemer protocols.

#### Controlled mutations

First, we provide a small simulation to illustrate a scenario similar to the motivational example described earlier. We simulate a string *s* of 100 random nucleotides and a string *t* derived from simulating mutations every 15th position in *s*. Insertions, deletions, and substitutions are chosen at random with equal probability of 1/3 each. We simulate *s* and *t* ten times to illustrate the variability in matches for the strobemers between simulations. The start positions of matching strobemers are shown in Fig. 2 under two different parametrizations for minstrobes and randstrobes. We note that we would not obtain any matches for k-mers of 15nt or larger in this scenario, and furthermore, no matches for spaced k-mers if the mutations were indels. Minstrobes, while more effective than k-mers in this scenario, fail to produce matches between many of the mutations for the (2,9,10,20) parametrization and for some with the (3,6,10,20) parametrization. We observe that randstrobes produce matches in all ten experiments under both parametrizations and provide a more random match distribution across the string than minstrobes. Hybridstrobes has a match performance in between minstrobes and randstrobes.

**Fig. 2.**
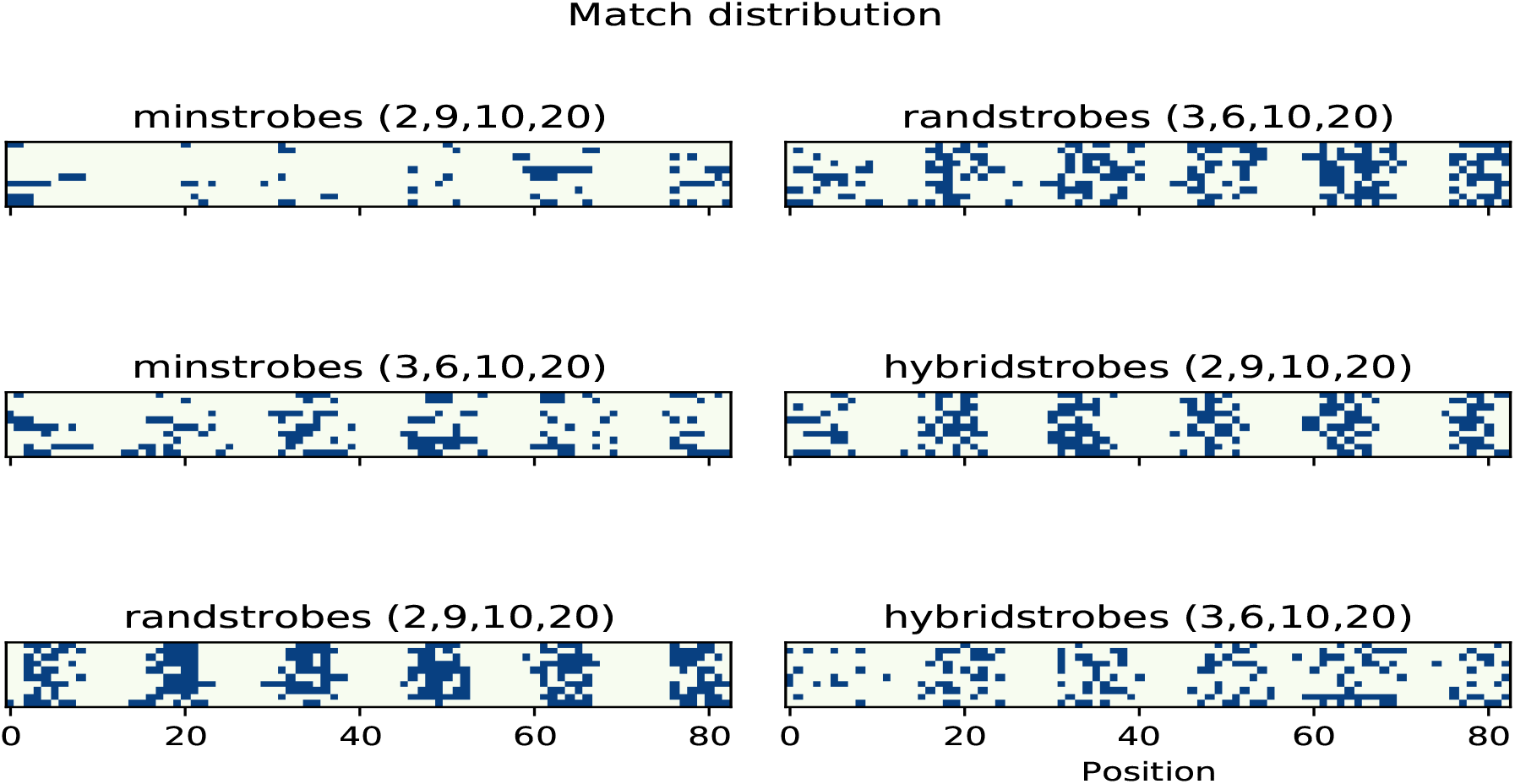
An example of strobemer matches for minstrobes, randstrobes and hybridstrobes with two different parametrizations each (separate panels). Each panel shows matches between a string *s* of 100nt and a string *t* derived from simulating mutations every 15th position in *s*. Indels and substitutions are chosen at random with equal probability. The matches are plotted with respect to the positions in *s* on the 83 possible matching positions (x-axis). Each row in a panel corresponds to a separate simulation.

To better quantify the performance in this scenario, we increase the size of our controlled experiment. We simulate a string of length 10,000nt and construct a second string by generating an insertion, deletion, or substitution with a probability of 1/3 each, every 20 nucleotides. We then simulate k-mers with size 30, spaced-dense with *k* = 30, and *L* = 45, spaced-sparse with *k* = 30, *L* = 90, and strobemers with parameters (2,15,25,50) and (3,10,25,50) so that all protocols have the same sampled subsequence length, and compare the number of matches, coverage, average island size, and island E-size under the different protocols (table 1). We repeat the experiment 1000 times to alleviate sample variance. For the spaced k-mer protocols, fixed positions are resimulated in each experiment. We refer to this as the SIM-C experiment (for simulation controlled). The spaced k-mer protocols offer an advantage over k-mers as they will match over some of the mutations that are substitutions. Particularly, the spaced-dense protocol that has a lower window size than space-sparse protocol and is therefore less affected by surrounding mutations. However, this particular experiment highlights the advantage of strobemers, which frequently produce matches between most or all mutations. Furthermore, the experiment shows the difference in performance between minstrobes, hybridstrobes, and randstrobes and their different parametrizations. The randstrobe protocols’ matches cover the largest fraction of the sequences, and they also have the smallest average and expected island size (table 1). In this experiment, the randstrobe of order 3 produces the most favorable sequence matching result. The hybridstrobe protocols have a match performance close to that of the randstrobe protocols across the four metrics.

**Table 1.**
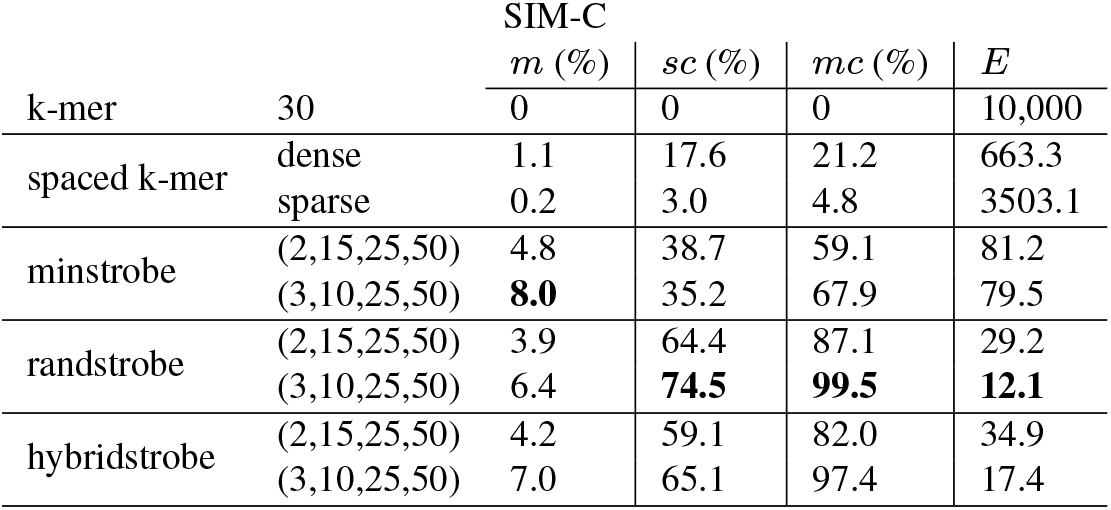
Statistics of the number of matches (*m*) as a percentage of the total extracted subsequences for the protocol, the sequence coverage (*sc*) and match coverage (*mc*) as a percentage of the total sequence length, and the expected island size (*E*) for the SIM-C dataset which has evenly spaced mutations with distance 20nt. The second column shows the parameters to the protocols.

#### Random mutations

In our second experiment, we simulate a string of length 10,000nt and construct a second string by generating insertions, deletions, or substitutions with equal probability of 1/3 each across the string with mutation rate *μ* ∈ 0.01,0.05,0.1. This means that the positions for the mutations are randomly distributed. Each such simulation is replicated 1000 times to alleviate sample variation. We refer to this as the SIM-R experiment (for simulation random). In this scenario, spaced k-mer protocols perform worse than k-mers, with fewer matches, lower match coverage, and larger expected island size (table 2). We observe that k-mers has the highest fraction of matches in all experiments. This is because matches produced by k-mers cluster optimally tight (1 nucleotide offset) between neighboring mutations at a distance larger than *k*. The minstrobe protocols under the two parametrizations have roughly the same performance as k-mers with higher match coverage and smaller expected island size but a lower fraction of matches and sequence coverage. The randstrobe protocols are also in this scenario significantly better at distributing matches across the sequences compared to all the other protocols. The randstrobe protocols have a substantially higher sequence coverage and match coverage and smaller expected island size under both parametrizations, which are all important aspects of sequence matching. Hybridstrobes produce results that are relatively close to the performance of randstrobes across the four matching metrics, making them a compelling alternative for sequence matching due to their fast construction time. We also show the full distribution of island sizes for mutation the different mutation rates (Fig. 3 and Fig. E.1) for a subset of the protocols, which illustrates the general trend in island sizes. For example, for a mutation rate of 0.1, we observe that the randstrobe protocols have roughly 1,000nt as the largest island size in our simulations, while k-mers have about 2,000nt (Fig. 3).

**Fig. 3.**
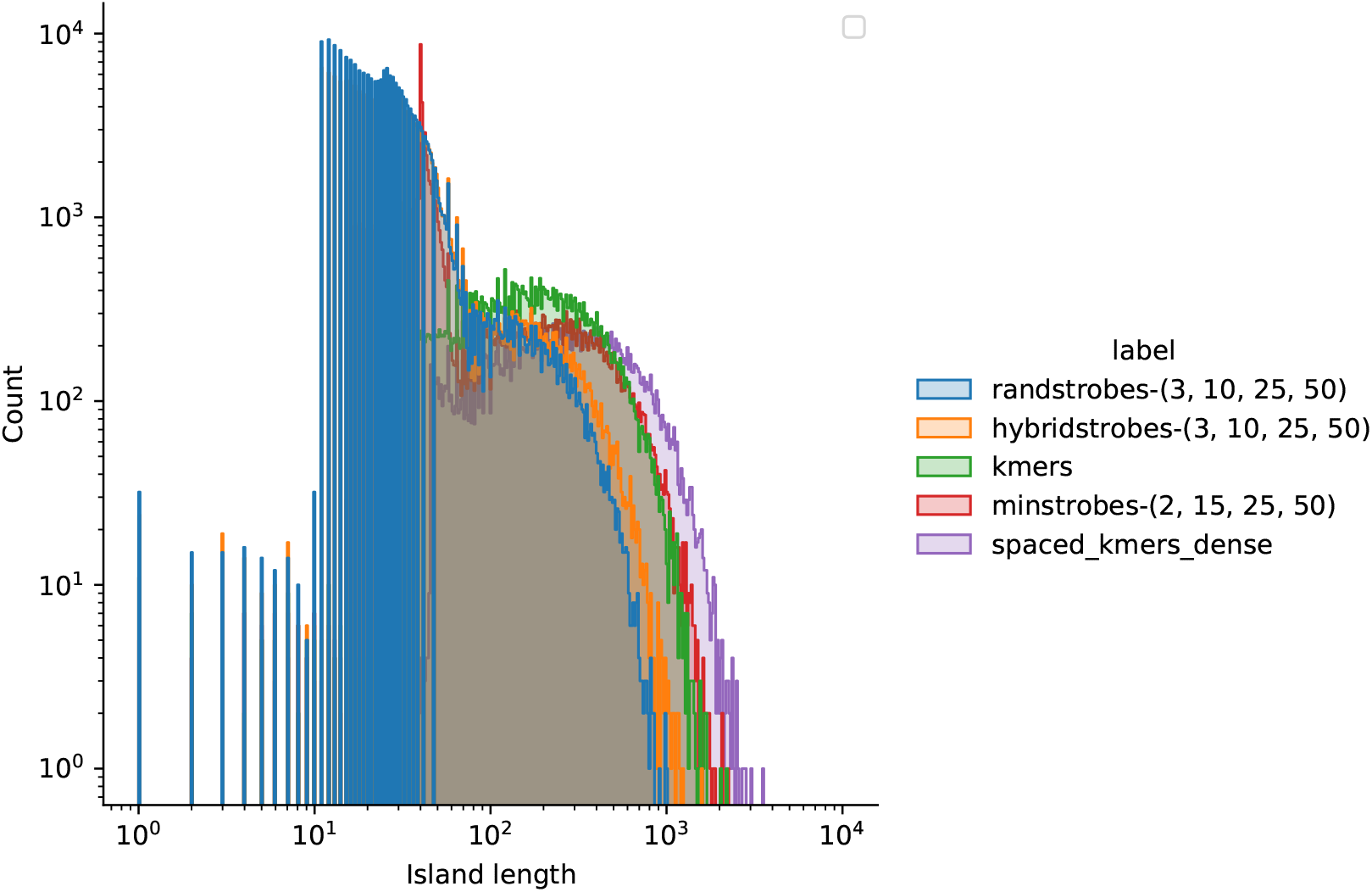
Histogram of island lengths for the SIM-R experiment with mutation rate 0.1.

**Table 2.** Match statistics under different sampling protocols under mutations rates of 0.01, 0.05, 0.1. Here, *m* denotes the number of matches as a percentage of the total number of extracted subsequences for the protocol, *sc* (sequence coverage) and *mc* (match coverage) is shown as the percentage of the total sequence length, and *E* is the expected island size.

|  |  | SIM-R |  |  |  | 0.01 |  |  |  | 0.05 |  |  |  |
| --- | --- | --- | --- | --- | --- | --- | --- | --- | --- | --- | --- | --- | --- |
|  |  | 0.1 |  |  |  |  |  |  |  |  |  |  |  |
| | | $m$ | $sc$ | $mc$ | $E$ | $m$ | $sc$ | $mc$ | $E$ | $m$ | $sc$ | $mc$ | $E$ |
| k-mer | 30 | **74.5** | 95.9 | 95.9 | 7.9 | **22.4** | 54.7 | 54.7 | 79.2 | **4.7** | 18.1 | 18.1 | 344.9 |
| spaced k-mer | dense | 67.6 | 95.6 | 96.2 | 9.7 | 13.8 | 50.9 | 53.9 | 120.7 | 1.8 | 14.1 | 16.1 | 570.1 |
|  | sparse | 50.5 | 87.8 | 89.7 | 44.4 | 3.5 | 21.4 | 26.7 | 640.7 | 0.1 | 2.1 | 3.6 | 4223.1 |
| minstrobe | (2,15,25,50) | 69.1 | 94.8 | 99.2 | 4.3 | 16.5 | 51.9 | 72.6 | 53.2 | 3.0 | 15.9 | 27.3 | 330.9 |
|  | (3,10,25,50) | 64.4 | 90.3 | 99.4 | 4.7 | 12.6 | 43.4 | 75.3 | 58.1 | 1.9 | 12.0 | 28.7 | 440.4 |
| randstrobe | (2,15,25,50) | 70.7 | 98.2 | 99.9 | 2.0 | 18.2 | 72.7 | 87.8 | 23.0 | 3.4 | 31.1 | 44.6 | 144.7 |
|  | (3,10,25,50) | 66.7 | **98.8** | **100.0** | **0.9** | 14.7 | **78.3** | **98.2** | **11.1** | 2.5 | **33.7** | **67.0** | **92.9** |
| hybridstrobe | (2,15,25,50) | 71.6 | 97.9 | 99.8 | 2.2 | 19.2 | 70.3 | 86.0 | 25.6 | 3.7 | 29.1 | 42.1 | 157.9 |
|  | (3,10,25,50) | 65.5 | 97.4 | 99.4 | 1.7 | 14.5 | 70.4 | 95.6 | 16.5 | 2.5 | 27.4 | 58.4 | 132.5 |

#### Thinning

K-mers, spaced k-mers and strobemers can all be thinned out using winnowing protocols such as minimizer schemes or syncmers (38). We compared the protocols when applying a minimizer protocol with thinning window size *w* =10 and 20 to both sequences in the SIM-R experiments. For k-mers and spaced k-mers, the thinning is performed by selecting the k-mer with the lowest hash value in a window of size *w*. For strobemers, the thinning is performed by selecting the the first strobe with the lowest hash value. This strobe will be selected to form the complete strobemer. In case of ties in hash values, the first k-mer (strobe) is selected.

In this scenario, the relative improvement of strobemers compared to k-mers decreases as *w* increases. For *w =* 10, randstrobes has a better sequence coverage, match coverage, and expected island size than all other protocols across mutation rates (table 3). With *w* = 20, k-mers produce the best sequence coverage across protocols, while randstrobes produce the best match coverage across protocols. Expected island size is better for randstrobes for mutation rates 0.01 and 0.05, but worse for mutation rate of 0.1. Hybridstrobes follows the performance of randstrobes closely in all experiments. Our experiments indicate that the relative increase in performance that strobemers have over k-mers decrease the more they are subsampled under the thinning protocol considered here.

**Table 3.**
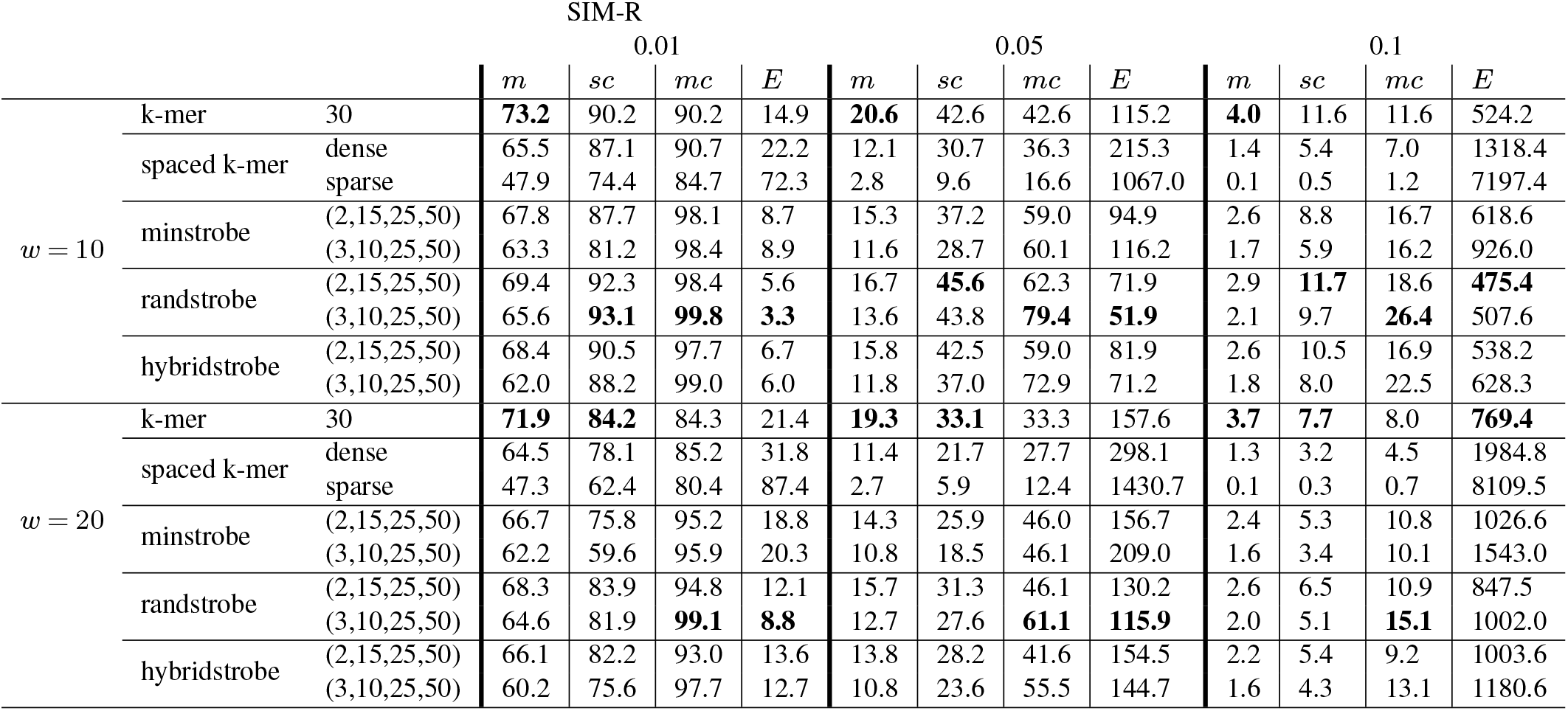
Match statistics under different sampling protocols under mutations rates of 0.01, 0.05, 0.1 using minimizer thinning with *w* = 10, and *w* = 20. Here, *m* denotes the number of matches as a percentage of the total number of extracted subsequences for the protocol, *sc* (sequence coverage) and *mc* (match coverage) is shown as the percentage of the total sequence length, and *E* is the expected island size.

### Strobemer vs k-mer uniqueness

We also want to compare the confidence or uniqueness of a match. Strobemers offer more match flexibility, as they can preserve a match with indels in the sampled region. We refer to the ability for a protocol to match over indels as *flexible-position* protocols), contrary to k-mers and spaced k-mers (referred to as *fixed-position* protocols). It is reasonable to assume that for the same size *k* of extracted subsequence, the strobemer protocols will have lower uniqueness (precision) than k-mers and spaced k-mers due to the flexible-position feature. We study the uniqueness in matches by computing the percentage of unique k-mers, spaced k-mers, and strobemers on the three largest human chromosomes (Fig. 4). Similarly to the SIM-C and SIM-R experiments, for a k-mer size of *k*, we parametrize the strobemer protocols with (*n,k/n*, 25,50) for *n* = 2,3 in order to have the same subsequence lengths. Similarly, the spaced k-mers are parametrized by *L* = 1.5*k* and *L* = 3*k* and the positions are simulated as in previous experiments.

**Fig. 4.**
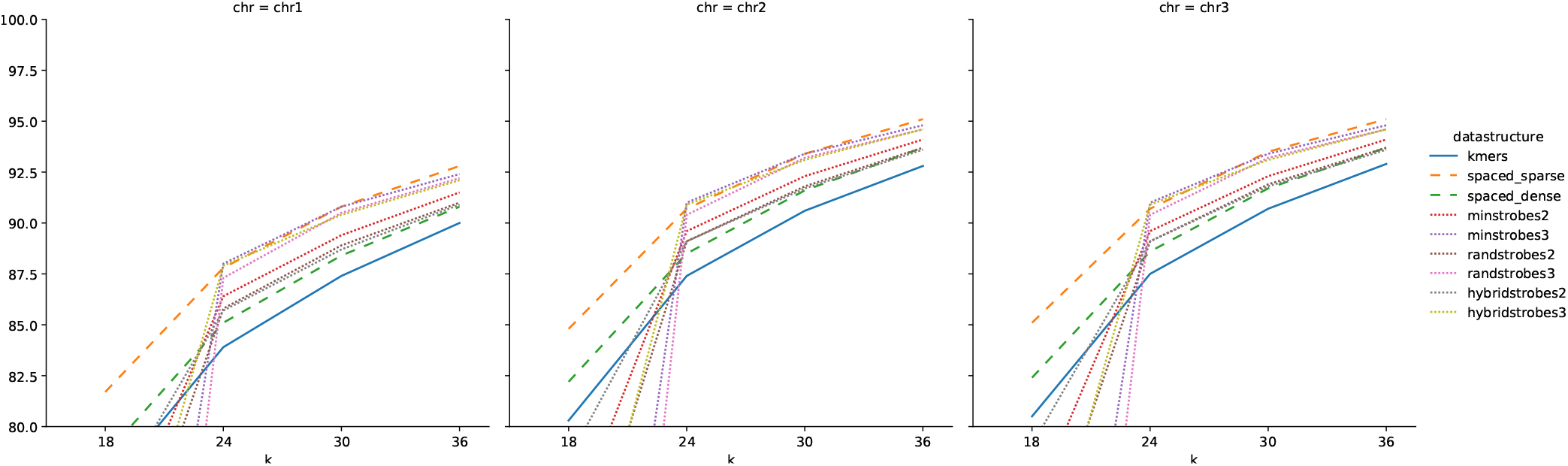
The percent of unique k-mers, spaced k-mers, minstrobes and randstrobes (y-axis) on the three largest chromosomes (chr1-chr3) of the human genome for various sequence lengths of *k* (x-axis). Each panel shows a separate chromosome. For a given *k* in the plot, strobemers with *n* = 2 are computed with parameters (2, *k*/2,50) and strobemers with *n* = 3 are computed with parameters (2, *k*/3,25) so that the number of extracted nucleotides between the five methods are the same. Y-axes have been cut at 80% for illustration. The values for minstrobes and randstrobes with parameters (3,6,25) are below 50% on all the three chromosomes. The value for minstrobes and randstrobes with parameters (2,9,50) are below 70% on all the three chromosomes.

We observe that for the three fixed-position protocols, a larger span in which positions are sampled helps subsequence uniqueness. The spaced-sparse has the highest uniqueness across all the three chromosomes, followed by spaced-dense and finally the k-mers.

Contrary to our intuition, the strobemer strobemers offer a higher uniqueness than k-mers for *k* ≥ 24 (Fig. 4), which may be due to the larger sampling window span as we observed for the spaced k-mers. The Out of the strobemer protocols evaluated here, strobemers of order 3 produce the highest percentage of unique matches for reasonably large subsequence lengths (*k* ≥ 24). There is no substantial difference between the strobemer protocols of the same order. However, for *k* = 18, the strobemer protocols will be parametrized by (2,9,25,50) and (3,6,25,25), which with the flexible-position sampling appear too small to guarantee reasonable uniqueness on the largest human chromosomes.

### Proof of concept sequence mapper

As demonstrated in our simulated experiments, spaced k-mers perform sub-optimally to k-mers and strobemers when indels are present. Therefore, we further compared k-mers to strobemers using synthetic and biological data with indels. We implemented a proof-of-concept tool StrobeMap. StrobeMap implements sequence similarity search with k-mers and strobemers of order 2 and 3. The output of StrobeMap is a tab separated values file (TSV) file with mapping information on the same format as MUMmer (48). However, instead of producing maximal exact matches (MEMs) or maximal unique matches (MUMs) between a query and a reference sequence, StrobeMap outputs what we refer to as Non-overlapping Approximately Matching (NAM) regions based on matches from the strobemer or k-mer protocol. The NAMs are produced by matches that overlap both on the query and reference, details on how NAMs are produced are found in appendix B.

As sequence mapping is often used as a preprocessing step to performing alignment or clustering, we use metrics valuable to candidate filtering to evaluate the methods. We measured the number of NAMs generated, the total match coverage produced by the NAMs, and the average normalized NAM length, which is the length of the NAM divided by either the length of the reference or the query depending on the mapping context. In order to achieve high accuracy and efficient sequence similarity searches, it is important that a mapping step produce few but long matches that cover a large portion of the query and/or the reference. Few matches will reduce time to post-cluster matches, reduce disk space (if matches are stored), while long contiguous matches will improve the decision on whether a candidate matching region should be aligned or not. We mapped ONT cDNA and DNA reads with 7% and 17% median error rate both to reference sequences and to the reads themselves. We also studied whole genome mapping of two *E. coli* genomes under some different settings. The details of the data and experiments are found in appendix B.

We first mapped cDNA reads (queries) to SIRVs (references) using k-mers and strobemers with subsequence size of 30 where strobemers were parametrized as (2,15,20,70) and (3,10,20,70). Randstrobes produce the highest match coverage to references (Fig. 5A), lowest number of matches (Fig. 5B)) and highest normalized NAM lengths (Fig. E.2). On this dataset, randstrobes are favourable to all other protocols when it comes to sequence matching, closely followed by hybridstrobes. Many of the NAMs that the randstrobes produce cover the full or near full SIRV reference (Fig. E.2). We observe the same trend when we compare the ability match reads to each other from the same SIRV (Fig. E.3). However, all the protocols produce a lower coverage and normalized match length due to the lower sequence identity.

**Fig. 5.**
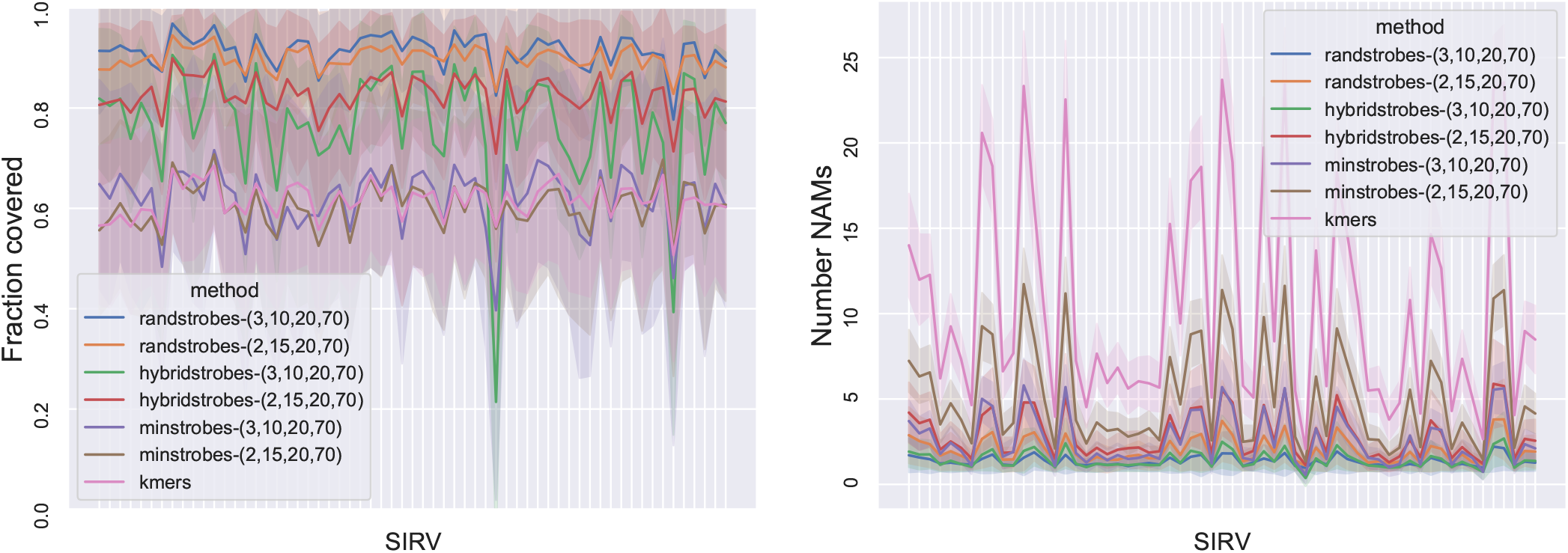
Comparison between strobemers and k-mers when matching ONT cDNA reads (7.0% median error rate) to 61 unique *Spike-in RNA Variants* (SIRV) reference sequences. Each SIRV corresponds to a tick on the x-axis. Panel **A** shows total fraction of the SIRV covered by NAMs from reads (y-axis). Panel **B** shows the number of NAMs (y-axis) between a read and the SIRV. The line shows the mean and the shaded area displays the standard deviation of the reads. A high NAM coverage and low number of NAMs means long contiguous matches and facilitates accurate and efficient sequence comparison.

When mapping genomic ONT *E. coli* reads to an *E. coli* genome, we measure how many NAMs the protocols generate and the fraction of the read that is covered by NAM matches coverage for the best mapping location. To get the best mapping location, we compute the longest collinear chain of NAMs to the genome. We count only the coverage of the longest collinear chain of NAMs to avoid overestimating coverage from additional matches (experiment details in appendix E). We compared k-mers of length 30 to hybridstrobes with parameters (3,10,10,100). The NAMs produced by hybridstrobes cover much more of the read (Fig. 6A) and are much fewer (Fig. 6B). We also mapped the reads against themselves and, similarly to mapping to the genome, we computed the total number of NAMs as well as the coverage of the longest collinear chain. This means that the coverage is only calculated for the largest overlap to another read. While we do not have the ground truth overlap values, large overlaps between pairs of longest overlapping reads are expected as the reads have a 3.65x coverage of the genome. Similarly to when we mapped the reads to the genome, we observe that hybridstrobes produce higher NAM coverage (Fig. 6C) and fewer NAMs (Fig. 6D).

**Fig. 6.**
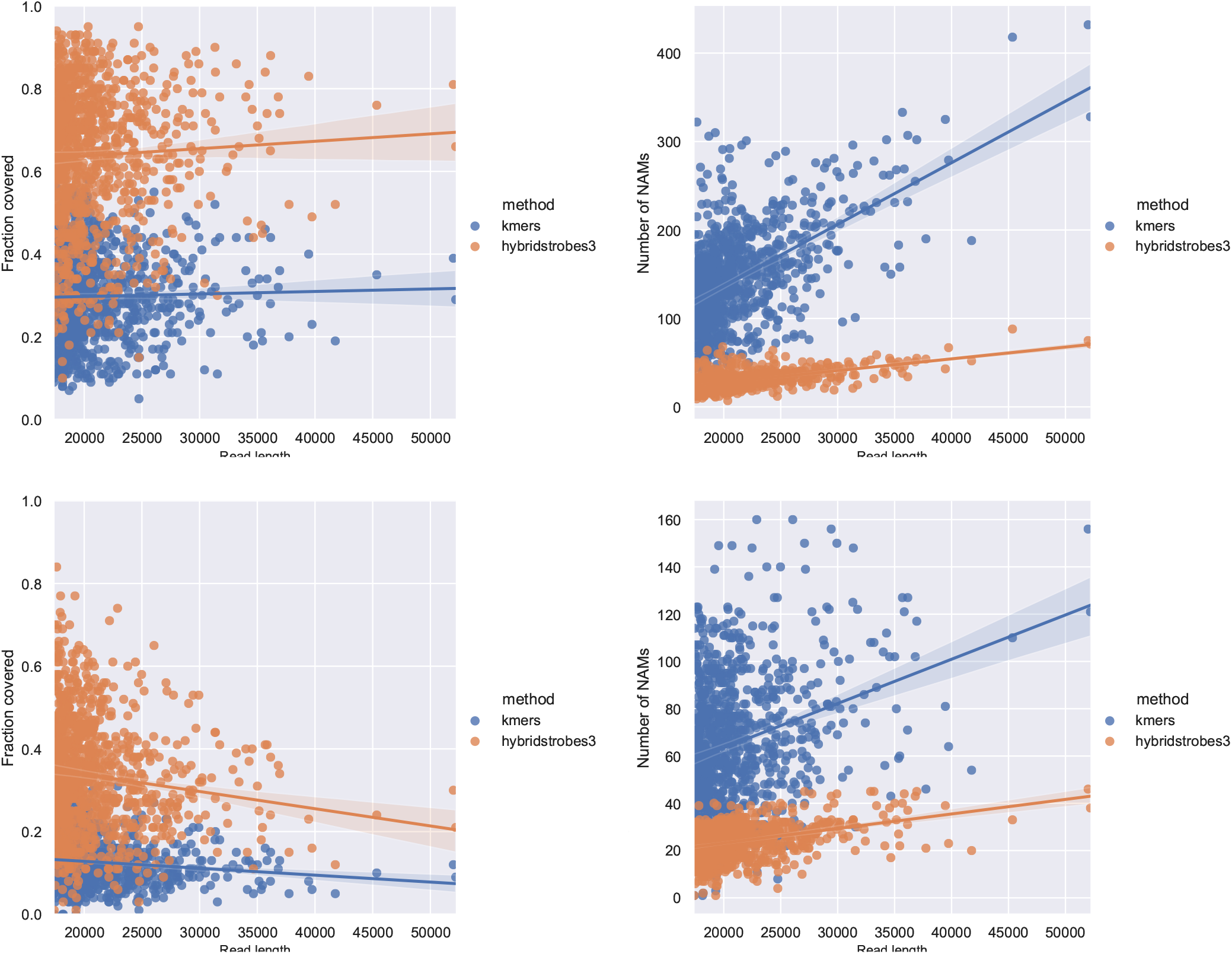
Comparison between hybridstrobes and k-mers when mapping genomic ONT reads for reads of different lengths (x-axis). Panel **A** and **B** shows read mapping results when mapping reads to the genome, and **C** and **D** when mapping reads to themselves. Panel **A** shows total fraction of the read covered by NAMs in the optimal colinear chaining solution to the genome (y-axis). Panel **B** shows the total number of NAMs (y-axis) between a read and the genome. Panel **C** shows the total fraction of the read covered by NAMs to the longest overlapping read, inferred from the optimal solution of a colinear chaining (y-axis). Panel **B** shows the total number of NAMs (y-axis) generated for the read. The line shows the mean and the shaded area displays a 95% confidence interval of the mean estimate. A high NAM coverage and low number of NAMs means long contiguous matches and facilitates accurate and efficient sequence comparison.

Finally, we measured the number of NAMs produced when we aligned two *E. coli* genomes to each other using k-mers of length 30 and hybridstrobes parameterized by (2,15,20,120), and (3,10,20,120). The k-mers produce 19,465 NAMs while hybridstrobes of order 2 and 3 produce 10,290 and 4,654 NAMs respectively. Fig. 7 shows MUMmer dotplots of the NAMs on the two *E. coli* genomes for the three mappings. Hybridstrobes produce long contiguous NAMs of similar regions and, with the parametrization here, avoids many of the smaller local matches. If one desires to also retain the smaller local hits, one can reduce the window sizes and *w_min_*. With hybridstrobes parametrized by (2,15,1,70), we retain most of the local hits with a total of 19,483 NAMs but still retaining the large contiguous matches (Fig. E.4A). The reason that the hybridstrobes produce slightly more NAMs compared to k-mers in this scenario is that *w_min_* is set so that the strobes can overlap in this example, producing smaller local hits. We also created hybridstrobes of order 3 with the parameters (3,10,20,120) and with minimizer thinning (*w* = 20) and observed similar long contiguous NAMs (Fig. E.4B), with a total of 3,213 NAMs produced. Details of the experiments are found in in appendix D.

**Fig. 7.**
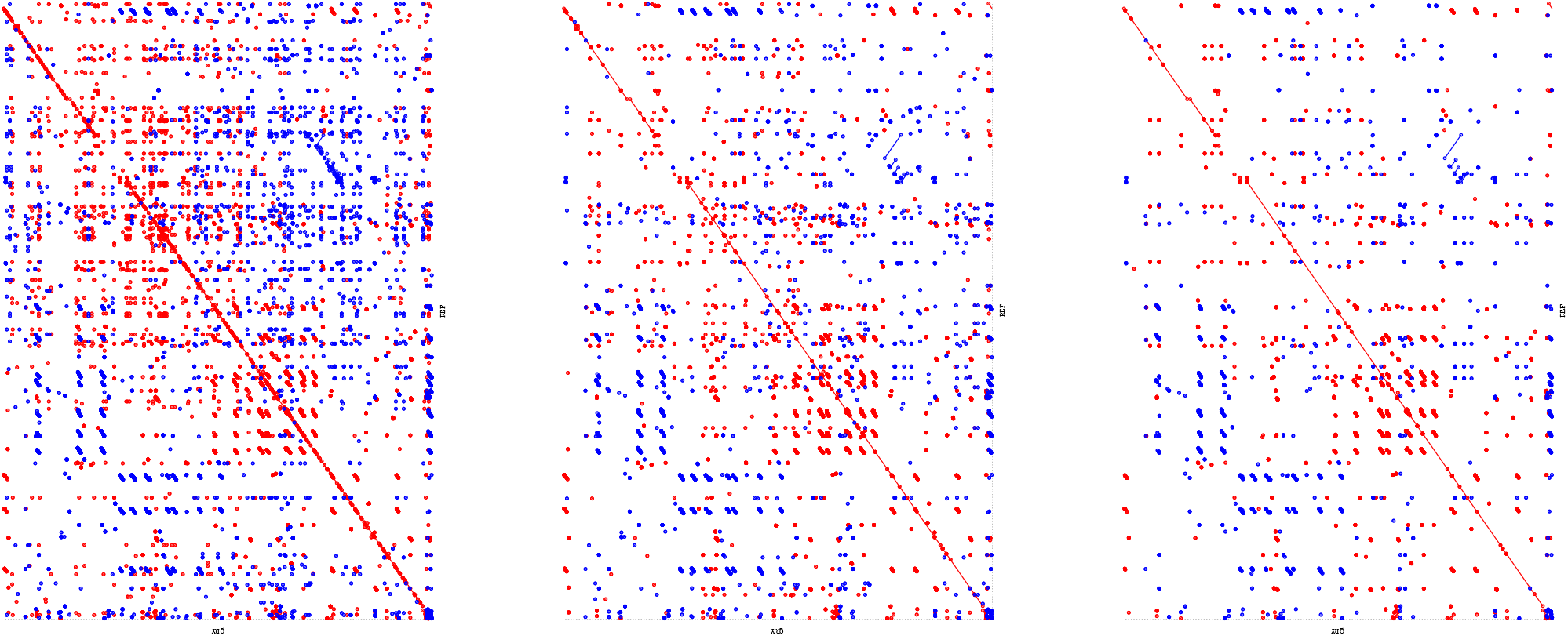
Dotplots of mapping two different *E. coli* genomes to eachother using (**A**) kmers of size 30, (**B**) hybridstrobes parametrized by (2,15,20,120), and (**C**) hybridstrobes parametrized by (3,10,20,120).

### Time and memory usage

For minstrobes and hybridstrobes, we only need to store queues with *w_max_* – *w_min_* + 1 hash values and the current minimum hash value in the queue. For randstrobes, the minimum of a hash value is computed from a window of size *w_max_* – *w_min_* + 1. The window size is negligible to sequence size for the window sizes investigated here. Therefore, the memory to construct strobemers is not significantly more memory intense than constructing k-mers.

Furthermore, strobemers take as much memory as k-mers to store for the same sequence length *k* and the start positions of k-mers or strobemers also require the same amount of memory. However, if one desires to store the positions of the other strobes, this could be done by storing offsets to the previous strobemer. For the window lengths investigated here, 8 bits per strobemer would suffice, with the possibility use less bits for smaller windows if only the offset to the start of the window is stored.

As for runtime, we compared the relative runtime of computing k-mers compared to strobemers using the construction described in the implementation section for different k-mer and window sizes (details of experiment in appendix C). K-mers are the fastest to compute. Randstrobes have the slowest relative runtime compared to k-mers, where the relative increase in computation time depends on the window size (table 3). Both minstrobes and hybridstrobes have comparable relative construction times to k-mers (table 3), making hybridstrobes, with their beneficial sequence match metrics, the most attractive protocol out of the strobemers.

However, we also show that the implementation and the programming language have substantial influence on the performance. We ran the same implementation under two different implementations of Python (Python 3.8 table 3 and pypy3 table 4) and observed drastic difference in the relative efficiency of computing strobemers (details in appendix C). Implementing strobemer protocols in a compiled programming language with arrays may decrease the relative construction time compared to k-mers (particularly for randstrobes; details in appendix C). Also, using single instruction multiple data (SIMD) implementations as is commonly used in bioinformatics (e.g., (49, 50)) may further improve relative construction time (see discussion in appendix C).

**Table 4.**
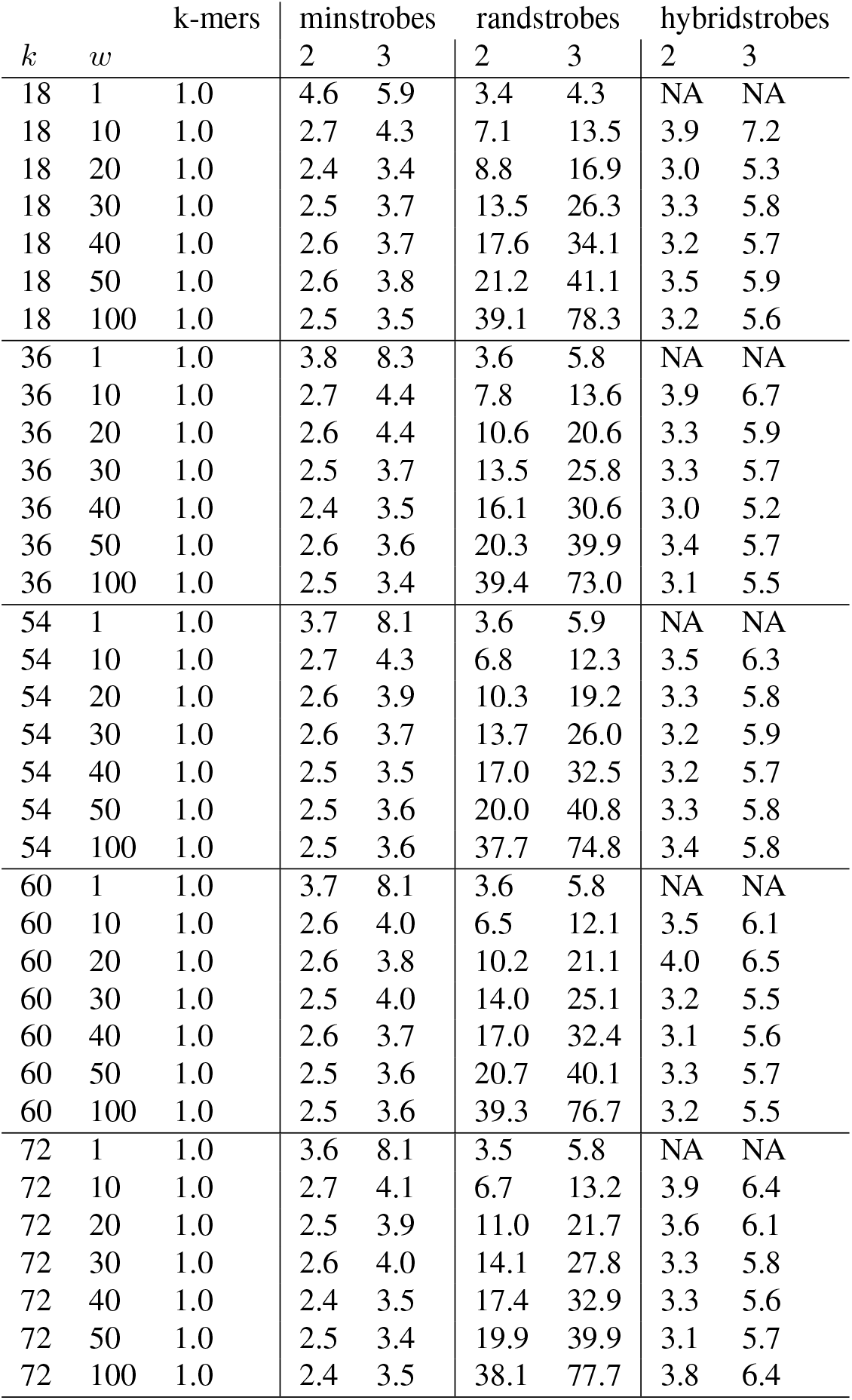
Relative time to compute k-mers compared to strobemers of order 2 and 3 using python v3.8 for various subsequence sizes (*k*) and window sizes (*w* = *w_max_* – *w_min_* + 1). The computation time is normalized with the time to compute k-mers. Hybrid strobes are not defined for window sizes smaller than *x* (the number of partitions of each window), which we here have set to 3.

When benchmarking memory consumption and runtime of our proof-of-concept tool StrobeMap on the *E. coli* data, we observed similar memory requirements for k-mers and hybridstrobes, while using hybridstrobes of order 2 is about twice as slow as k-mers, and hybridstrobes of order 3 is about a three to four times as slow in relative runtime. The benchmarks are given for reference but may not be representative of neither runtime, nor memory requirement of optimized implementations in a compiled programming language. The details of runtime and memory usage are found in appendix D. In addition, downstream processing of matches (such as collinear chaining) may take longer for k-mers as they in general producing significantly more matches.

## Discussion

We have studied strobemers, an alternative sampling protocol to k-mers and spaced k-mers for sequence comparison. We have experimentally demonstrated that strobemers, particularly randstrobes and hybridstrobes, efficiently produce higher sequence coverage, match coverage, and lower gap size between matches under different mutation rates (table 1 and table 2). Strobemers also produce a higher number of unique matches (specificity) compared to k-mers for several commonly used sized of *k* (Fig. 4).

K-mers produce the highest number of matches in the SIM-R experiments, as k-mer matches cluster optimally tight between mutations at distance larger than *k*. However, the number of matches is not always helpful as matches may cluster due to local repeats. Randstrobes and hybridstrobes can offer more evenly distributed matches, higher match coverage, and higher uniqueness. These are features that are useful for several algorithms that require chains of matches between two sequences to be considered candidates for alignment or clustering, e.g., as in (31, 33).

To show the utility of strobemers, we implemented a proof-of-concept mapping tool, StrobeMap, that perform sequence mappings using both k-mers and strobemers. We demonstrated in several different scenarios such as mapping ONT cDNA and genomic reads to themselves or to reference transcripts or genomes that strobemers produce favourable sequence comparison metrics. Particularly, hybridstrobes offer a beneficial trade-off between construction time and the ability to produce long contiguous matches under various sequence matching contexts. Overall, the strobemers, show a promising data structure for algorithms that rely on sequence comparison.

Similarly to k-mers, minstrobes and randstrobes can be subsampled as minimizers (29), syncmers (38), or any other thinning protocol that can be applied to k-mers. We observed that the more thinned out the strobemer protocols are, the less advantage do they have over k-mers (table 3). An interesting future research direction would be to study whether specific thinning schemes are better suited for strobemers. Specifically, whether they can preserve the relative performance increase that are observed without thinning. By studying the mathematical properties of hashes and minimizers (36, 51), we may find a effective subsampling techniques of strobemers.

As for runtime performance randstrobes are slower to generate than k-mers and minstrobes. Hybridstrobes mixes the ideas from minstrobes and randstrobes and shows a runtime comparable to minstrobes while producing sequence matches almost as efficiently as randstrobes. Overall, we believe hybridstrobes may offer the best trade-off in performance and sequence comparison accuracy. However, by employing ideas like cyclic polynomial hash functions (52), we may come up with faster methods to generate strobemers.

### Future study of strobemers

#### Parameterization

While our study provides an experimental evaluation of strobemers under some commonly used values of *k* and mutation rates, the statistics of strobemers remains to be explored. In (18), the authors derived the mean and variance of islands for k-mers and the number of mutated k-mers under given mutation rate. If we can derive analytic expressions for strobemers, it may suggest us how to optimize parameters of the strobemer protocols under various mutation rates, which will be useful for similarity comparison algorithms. Even without analytic expressions, we can evaluate the sizes on strobes and windows suitable for various mutation rates. Also, we could relax the constraint of equal-size strobes and window sizes. As a start in this direction, we may derive more efficient parameter selection on window sizes by modeling the number of mutations after a certain number of nucleotides as a Poisson Process. Under such a model, the author hypothesizes that choosing larger window sizes downstream could be beneficial. This remains to be explored.

#### Construction, storing and queries

There are several aspects of construction, indexing, and storage of strobemers that could be explored. One such direction is to store and query the positions of the other strobes efficiently, as they give extra information about the coverage and span of matches across sequences for sequence similarity applications. Another application is to efficiently index the data sets for abundance and presence of strobemers (53). For such applications, minstrobes may be advantageous due to the more frequently shared minimizers between the strobes. Finally, the possibility of decreasing practical runtime for constructing randstrobes remains to be explored.

#### Span-coverage for matching

Since strobemers are gapped sequences, it also motivates the study of match coverage and distribution of matches across regions (or positions) similarly to what has been done for gapped experimental protocols such as mate-pair or paired-end reads (54). For example, one could compute the span-coverage of matches at positions or over regions to estimate the sequence similarity in matching regions or the confidence for further downstream processing.

#### Generalization of strobemers

We can view the process of extracting a k-mer or a spaced k-mer at position *i* in a string *s* as applying a function *f*(*i,k,s*) on *s*. Similarly, the process of extracting a strobemer from *s* can be viewed as applying the higher-order function *f*′(*i,k,s,h*) on *s* where *h* is either some hash function or hash strategy (e.g., iterative and conditionally dependent as in randstrobes). We demonstrated that applying *f*′ on *s* is equally or more efficient than *f* for sequence matching for three different functions *h* (minstrobes, randstrobes, and hybridstrobes), which poses the following question. Can we further improve sampling protocols for sequence matching by designing *h* differently?

## Conclusions

We have presented strobemers as an alternative to k-mers and spaced k-mers for sequence comparison. Strobemers, particularly randstrobes and hybridstrobes, offer a more evenly distributed set of matches across sequences compared to k-mers and spaced k-mers, are less sensitive to the distribution of mutations across sequences, and produce a higher match coverage under several parameterizations. We also showed that strobemers can offer higher match uniqueness compared to k-mers for several same subsequence lengths. These features are useful for algorithms that perform sequence matching. Strobemers are also easy to both construct and query, making it a compelling alternative to k-mers. We also demonstrated the utility of strobemers in several sequence comparison applications using synthetic and biological sequencing data. While we have empirically demonstrated the useful properties of strobemers, their statistical properties require further investigation.

## ACKNOWLEDGEMENTS

We thank Camille Marchet, Robert Harris, Rayan Chikhi, Paul Medvedev, Lior Pachter, Karel Břinda, Michael Hall, Michael Schatz, and Páll Melsted for their constructive comments and suggestions on an early draft of the manuscript. The computations were performed on resources provided by the Swedish National Infrastructure for Computing (SNIC) at Uppsala Multidisciplinary Center for Advanced Computational Science (UPPMAX).

## Supplementary Note A: Strobemers construction

**Algorithm 1:**
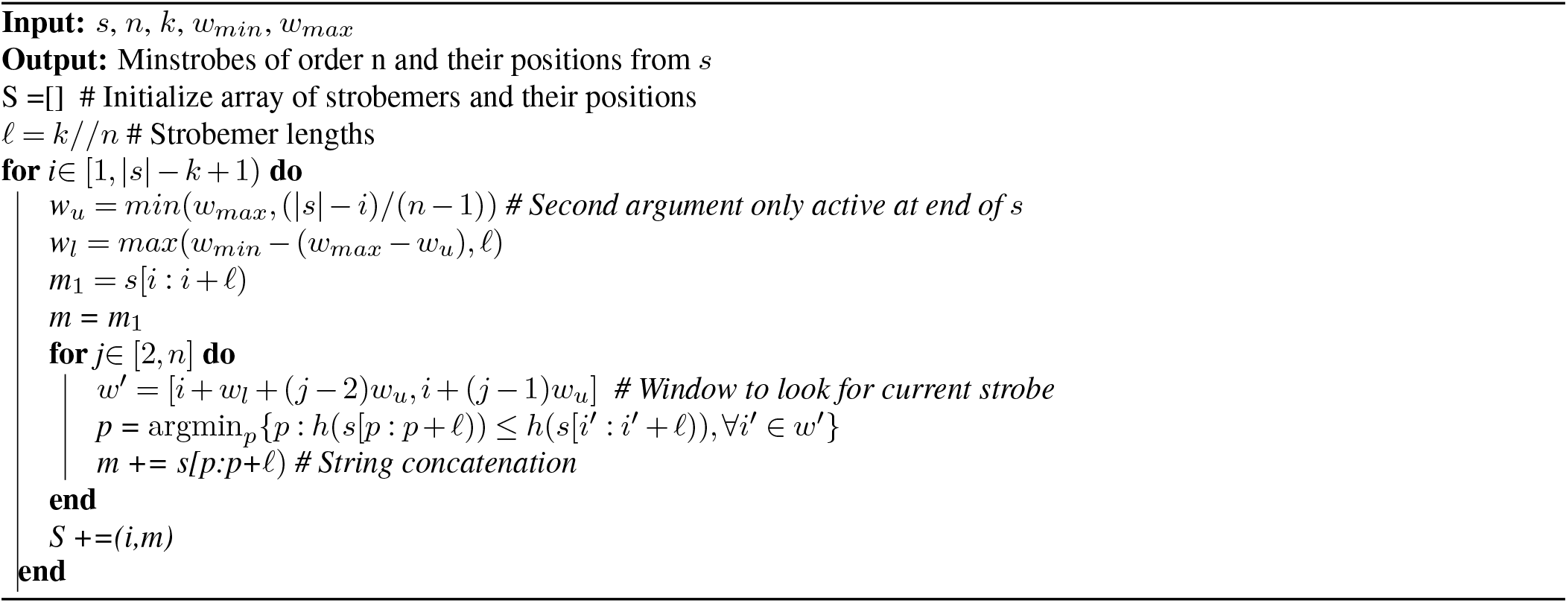
Minstrobes construction

**Algorithm 2:**
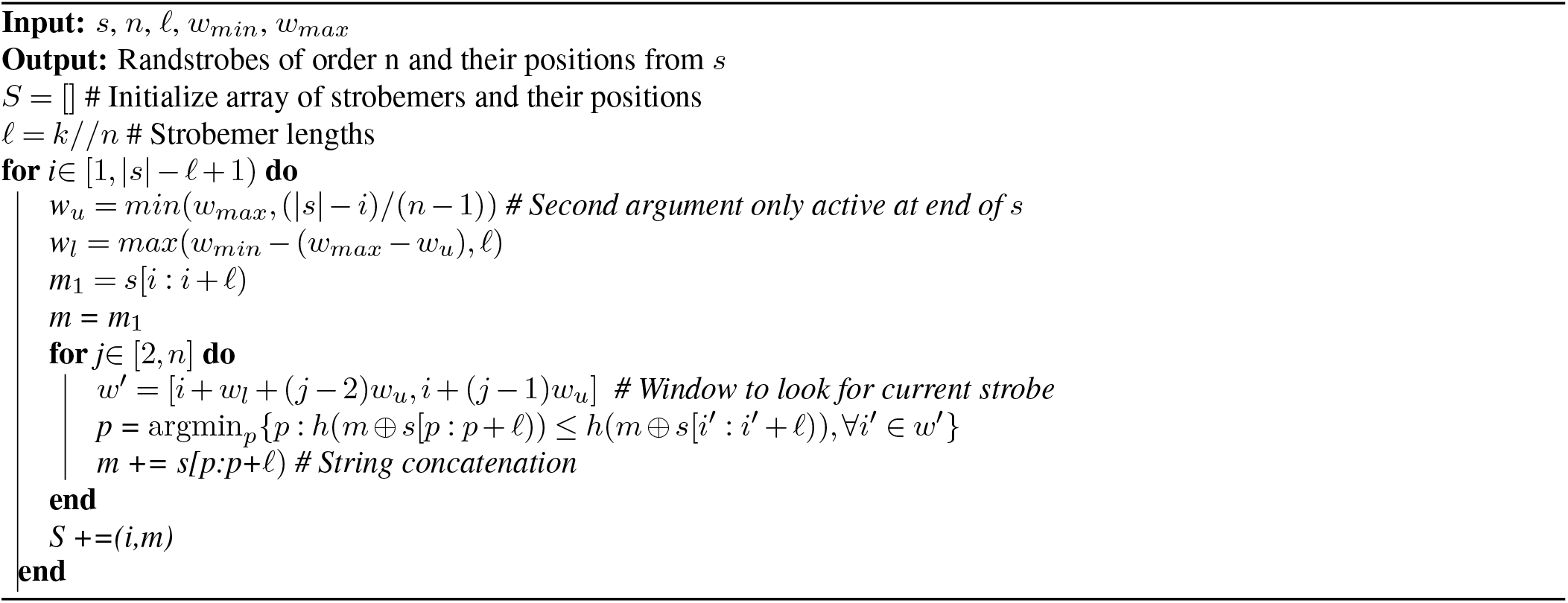
Randstrobes construction

**Algorithm 3:**
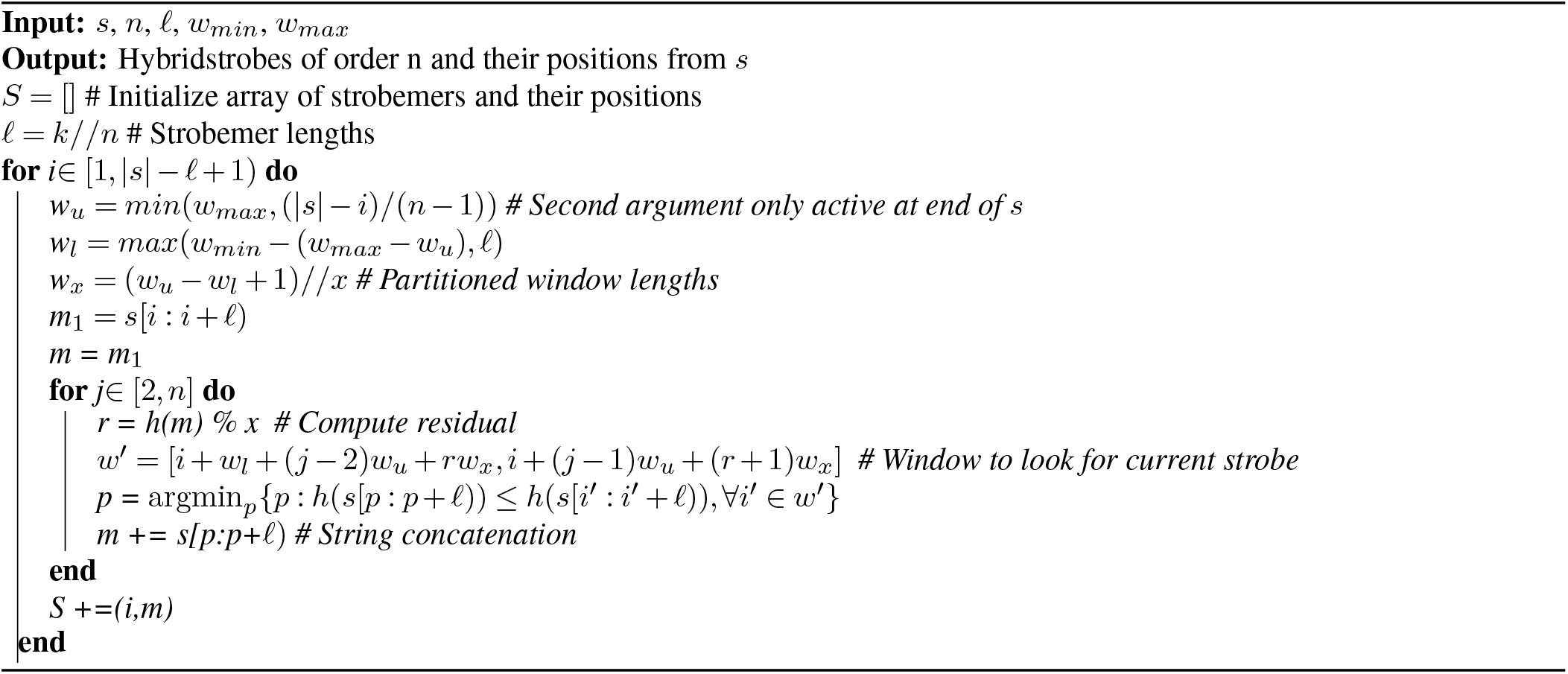
Hybridstrobes construction

## Supplementary Note B: Mapping analysis

### A. Constructing Non-overlapping Approximate Matches (NAMs)

The NAMs are produced as follows. Assume a query sequence *q* and a reference sequence *r*, and two strobemers *a* and *b* where the match of *a* spans positions [*q_i_,q_j_*] and [*r_i_,r_j_*] and the match of *b* spans [*q_i′_,q_j′_*] and [*r_i′_,r_j′_*] on *q* and *r*, respectively. If it holds that *q_i′_* ≤ *q_i′_* ≤ *q_j_* and *r_i_* ≤ *r_i′_* ≤ *r_j_* we say that they *overlap* where *a* precedes *b* and *b* supersedes *a*. A NAM spanning [*q*_1_,*q*_2_] och the query sequence and [*r*_1_, *r*_2_] on the reference is a chain of overlapping strobemers such that no other strobemer produce neither a preceding nor a superseding overlap with the NAM on both *q* and *r*. The definition of NAMs for k-mers is identical where a k-mer match, as opposed to a strobemer match, spans k consecutive positions. Note that a NAM is not the same as a MEM even in the case of k-mers as, e.g., a length difference in a homopolymer region will break a MEM but will not break a NAM formed from k-mers.

### B. SIRV reads to SIRV references

We downloaded SIRV ONT cDNA reads from ENA with accession number PRJEB34849, and SIRV references from https://www.lexogen.com/wp-content/uploads/2018/08/SIRV_Set1_Lot00141_Sequences_170612a-ZIP.zip. We preprocessed the cDNA reads using pychopper (https://github.com/nanoporetech/pychopper) to produce full-length cDNA sequences. We then aligned the full-length reads using minimap2 (55) to the references and subsampled 100 reads from each reference. For each SIRV we subsampled from the pool of reads that had a primary alignment to the SIRV that started and ended not more than ten nucleotides from the start and end of the SIRV, respectively. This was done to assure that in an ideal matching scenario, a NAM from a read to a SIRV could cover the entire SIRV.

### C. SIRV reads to each other

In this experiment, we took the 100 reads subsampled as described in the previous section, and mapped each of the 100 reads to the other 99 reads within the pool for each SIRV.

### D. *E. coli* genomes to themselves

We mapped the *E. coli* genome GCA_003018135.1 ASM301813v1 to the *E. coli* genome GCA_003018575.1 ASM301857v1 available at https://www.ncbi.nlm.nih.gov/genome/167?genome_assembly_id=368391.

We ran StrobeMap as follows

~~~
StrobeMap --queries GCF_003018135.1_ASM301813v1_genomic.fna
            --references GCF_003018575.1_ASM301857v1_genomic.fna
            --outfolder out/ --rev_comp
~~~

with the specific parameter

~~~
--k 30 --kmer_index
~~~

to produce k-mer NAMs and, for example,

~~~
--k 15 --strobe_w_min_offset 20 --strobe_w_max_offset 120 --n 2 [--w 20]
~~~

to produce hybridstrobes with parameters (2,15,20,120).

### E. *E. coli* reads to E.coli genome

We downloaded *E. coli* reads from Sequence read archive with Run ID SRR13893500. As the sample contains a fraction of reads from other bacteria, we selected the 1000 longest reads from the sample that aligned to the *E. coli* genome with more than 95% of the total read length. As aligned portion we computed the span between the first and last base that was aligned to the genome divided by the read length. This calculation excludes hard and softclipped ends but does consider eventual poorly aligned internal regions of the read. This produced 1,000 reads with a median length of 19,601nt where the longest read was 52,197nt and the shortest read was 17,360nt. The total length of the reads was 21,020,364 giving a coverage of 6.65x. To obtain the subsampled reads, we ran minimap2 as follows and a custom script available in the strobemer repository to select the reads from the SAM file.

~~~
minimap2 -ax map-ont --eqx GCF_003018135.1_ASM301813v1_genomic.fna \
                            SRR13893500.fastq > SRR13893500.sam
python select_longest_reads.py SRR13893500.fastq SRR13893500.sam \
                                1000 SRR13893500_1000_longest.fastq
~~~

We further estimated the read error rate from these reads by dividing the sum of substitutions and indels with the length of the aligned region to get a median error rate of 17.0%. We mapped the 1,000 reads using StrobMatch to the *E. coli* genome GCA_003018135.1 ASM301813v1 available at https://www.ncbi.nlm.nih.gov/genome/167?genome_assembly_id=368373 and measured the number of NAMs and the match coverage produced by the colinear chain of matches that covers the largest fraction of the reads. We count the coverage only for the colinear chains as smaller matches to other region of the genome may inflate the coverage of the read to the actual best aligned region. However, all matches to the genome contribute to the total number of matches, because this is an important efficiency metric to select candidate alignments.

## Supplementary Note C: Runtime analysis

We compared the relative runtime of computing k-mers compared to strobemers using the construction described in the implementation section. We used python v3.8 for the experiments. We simulated 1000 strings of length 100,000nt and computed the runtime to extract k-mers and strobemers under different subsequence sizes (18,36,54,60,72) and window sizes (1,10,20,30,40,50,100). The time to construct the strobemers is normalized with the time to construct k-mers. Runtime results are showed in table 4.

In general, randstobes are the slowest to construct. For randstrobes, the construction time increases with window size, where the construction time shows roughly a linear relationship with the window size. As expected, the construction time also increases with the number of strobes. Minstrobes and hybrid strobes relies on the implementation of queues in python. For these protocols, they have a better performance with larger window sizes, as the queues do not need to be updated as frequently. We note that minstrobes and hybridstrobes have a similar runtime performance.

This runtime comparison comes with many caveats. First, it depends on out implementation and data structures used. Secondly, it is further highly dependent on the programming language in which they are implemented. To illustrate this, we ran the exact same code using pypy v7.3.3-beta (table 5). In this scenario, randstrobes are now, in relative terms, more than twice as fast to compute compared to under python 3.8. Furthermore, minstrobes and hybridstrobes now suffer greatly for small window sizes where the queues frequently needs to be updated. The relative numbers between k-mers and strobemers will change under an other implementations and programming language. This illustrates that the it is perhaps more informative to investigate the time complexity for the protocols in this scenario, as programming languages may have different overheads and datastructure specific performances.

**Table 5.** Relative time to compute k-mers compared to strobemers of order 2 and 3 using pypy3 for various subsequennce sizes (*k*) and window sizes (*w* = *w_max_* – *w_min_* + 1). The computation time is normalized with the time to compute k-mers.

| $k$ | $w$ | k-mers | minstrobes | | randstrobes | | | |
| --- | --- | --- | --- | --- | --- | --- | --- | --- |
|  |  |  | 2 | 3 | 2 | 3 | 2 | 3 |
| 18 | 1 | 1.0 | 3.9 | 7.6 | 3.0 | 4.9 | NA | NA |
| 18 | 10 | 1.0 | 3.0 | 5.0 | 5.3 | 10.1 | 10.6 | 60.1 |
| 18 | 20 | 1.0 | 2.9 | 4.3 | 7.1 | 13.3 | 8.2 | 37.4 |
| 18 | 30 | 1.0 | 2.8 | 4.4 | 8.7 | 18.2 | 7.8 | 35.1 |
| 18 | 40 | 1.0 | 2.7 | 3.8 | 8.8 | 18.1 | 6.3 | 20.0 |
| 18 | 50 | 1.0 | 2.8 | 4.2 | 12.8 | 26.0 | 5.8 | 16.3 |
| 18 | 100 | 1.0 | 2.7 | 4.1 | 19.1 | 42.7 | 4.8 | 10.2 |
| 36 | 1 | 1.0 | 8.1 | 12.9 | 3.9 | 5.7 | NA | NA |
| 36 | 10 | 1.0 | 3.3 | 5.5 | 5.5 | 10.0 | 8.3 | 49.6 |
| 36 | 20 | 1.0 | 2.6 | 4.1 | 6.4 | 12.4 | 8.1 | 35.3 |
| 36 | 30 | 1.0 | 2.7 | 4.4 | 8.9 | 17.7 | 5.5 | 23.0 |
| 36 | 40 | 1.0 | 2.6 | 4.0 | 10.3 | 19.9 | 4.9 | 17.0 |
| 36 | 50 | 1.0 | 2.7 | 3.9 | 11.9 | 26.0 | 5.1 | 14.9 |
| 36 | 100 | 1.0 | 2.7 | 4.0 | 20.6 | 44.7 | 4.5 | 10.4 |
| 54 | 1 | 1.0 | 3.4 | 7.4 | 3.3 | 5.0 | NA | NA |
| 54 | 10 | 1.0 | 2.5 | 4.5 | 5.3 | 8.7 | 7.1 | 41.7 |
| 54 | 20 | 1.0 | 3.0 | 4.4 | 6.5 | 13.2 | 8.3 | 37.0 |
| 54 | 30 | 1.0 | 2.8 | 4.5 | 8.1 | 16.1 | 5.8 | 22.8 |
| 54 | 40 | 1.0 | 2.7 | 4.1 | 9.6 | 19.7 | 5.1 | 17.0 |
| 54 | 50 | 1.0 | 2.7 | 4.0 | 11.0 | 22.9 | 5.0 | 14.0 |
| 54 | 100 | 1.0 | 2.6 | 3.8 | 17.8 | 38.4 | 4.2 | 10.0 |
| 60 | 1 | 1.0 | 7.0 | 12.0 | 3.5 | 5.0 | NA | NA |
| 60 | 10 | 1.0 | 3.1 | 4.9 | 5.0 | 8.8 | 7.1 | 42.0 |
| 60 | 20 | 1.0 | 2.8 | 4.3 | 6.4 | 12.2 | 7.9 | 34.3 |
| 60 | 30 | 1.0 | 2.8 | 4.4 | 8.4 | 16.5 | 7.8 | 28.5 |
| 60 | 40 | 1.0 | 2.8 | 4.2 | 9.8 | 20.0 | 7.4 | 23.2 |
| 60 | 50 | 1.0 | 2.8 | 4.0 | 11.8 | 22.9 | 6.5 | 18.9 |
| 60 | 100 | 1.0 | 2.6 | 3.9 | 18.1 | 39.1 | 6.4 | 10.0 |
| 72 | 1 | 1.0 | 7.0 | 10.8 | 3.3 | 4.7 | NA | NA |
| 72 | 10 | 1.0 | 3.2 | 5.0 | 5.0 | 8.8 | 8.3 | 49.0 |
| 72 | 20 | 1.0 | 2.8 | 4.5 | 6.6 | 12.5 | 8.2 | 36.7 |
| 72 | 30 | 1.0 | 2.7 | 3.9 | 7.6 | 15.2 | 7.1 | 26.3 |
| 72 | 40 | 1.0 | 2.7 | 4.0 | 9.3 | 19.0 | 7.1 | 23.3 |
| 72 | 50 | 1.0 | 2.7 | 4.0 | 11.2 | 24.0 | 7.0 | 20.7 |
| 72 | 100 | 1.0 | 2.6 | 3.7 | 17.1 | 39.7 | 5.4 | 9.7 |

There are however several implementation tricks that one could perform to speed up construction time of, e.g., randstrobes. In our python implementation, we have used lists to store hash values which implemented as arrays of pointers to the integers and slow to iterate and compute minimum over. Using arrays in compiled languages could further speed up computation. I addition, it is possible that the computation of strobemers, particularly randstrobes, could be sped up using, e.g., single instruction multiple data (SIMD) implementations as is commonly used in bioinformatics (49, 50).

We excluded construction of spaced k-mers since they are, in our implementation, very time consuming to construct in python. For efficient construction of spaced k-mers, an array based compiled programming language should be used.

## Supplementary Note D: Memory requirement of StrobeMap

When benchmarking memory consumption used by our proof-of-concept tool StrobeMap, the peak memory usage for mapping the two *E. coli* genomes to each other was 2.88Gb, 2.60Gb, 2.93Gb, for k-mers, hybridstrobes of order 2, and hybridstrobes of order 3, respectively. The total time to produce the mapping was 34, 58, and 85 seconds for k-mers, hybridstrobes of order 2, and hybridstrobes of order 3, respectively. For the experiments mapping reads to the *E. coli* reference, the total time to produce the mappings was 44 and 179 seconds for the k-mers and hybridstrobes of order 3, respectively. The total memory consumption was 1.43Gb for k-mers and 0.97Gb for the hybridstrobes. For the experiments mapping *E. coli* reads to themselves, the total time to produce the mappings was 107 and 280 seconds for the k-mers and hybridstrobes of order 3, respectively. The total memory consumption was 3.48Gb for k-mers and 3.45Gb for the hybridstrobes. The large discrepancy in the memory consumption mapping reads to the *E. coli* genome is caused by much fewer NAMs produced by the hybridstrobes in combination with that StrobeMap stores in memory the NAMs for batches or reads only to invoke printing to file (and clearing matches) every thousand reads. In our experiments, using k-mers is fastest while hybridstrobes of order two took roughly twice as long and hybridstrobes of order 3 about 3-4 times as long.

## Supplementary Note E: Figures

**Fig. E.1.**
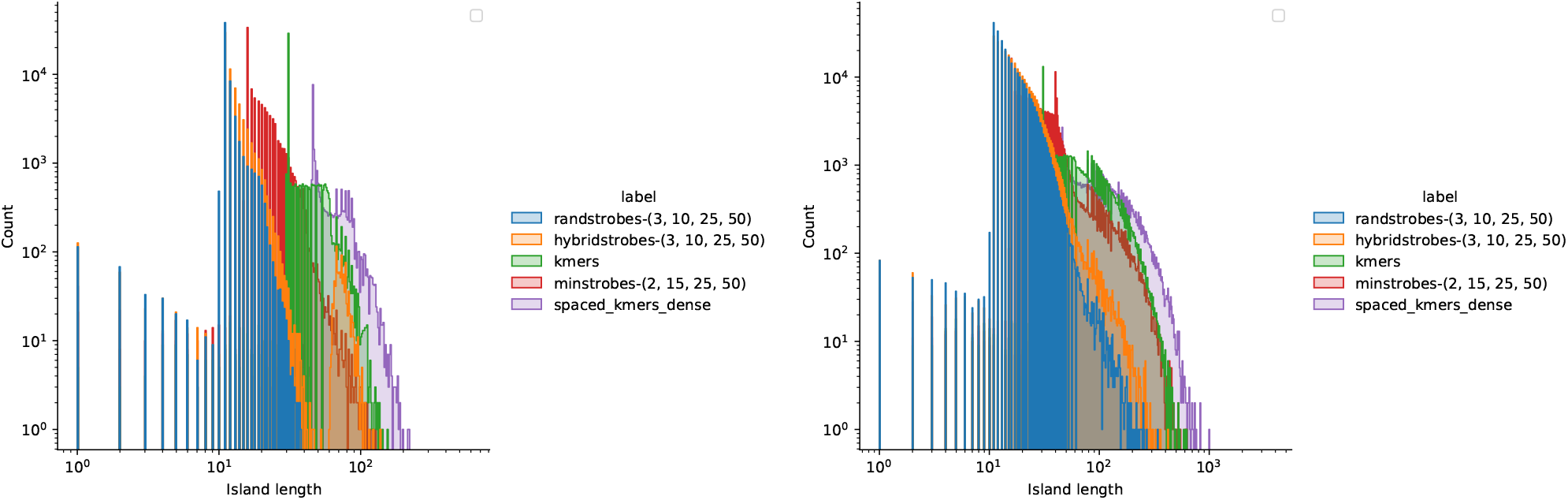
Histograms of island lengths for the SIM-R experiments for mutation rate 0.01 (a) and 0.1 (b).

**Fig. E.2.**
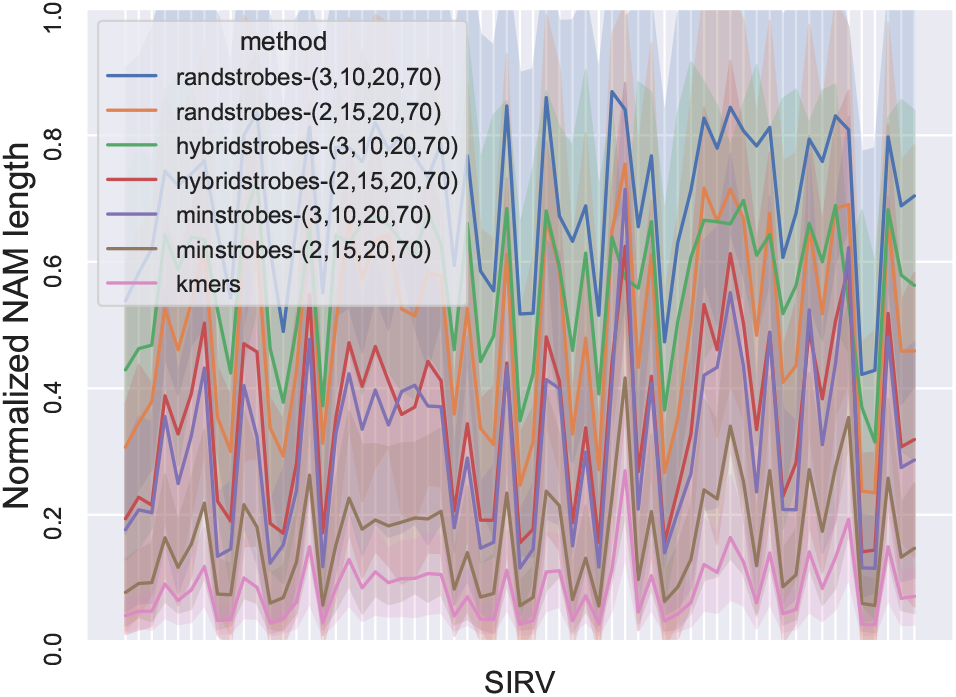
The plot shows the average normalized NAM length from all the NAM matches when matching ONT cDNA reads to 61 unique SIRV reference sequences. The normalized NAM length is the length of the NAM divided by the SIRV reference. Each tick on the x-axis corresponds to a SIRV. The line shows the mean and the shaded area displays the standard deviation of the reads. A high NAM coverage and low number of NAMs means long contiguous matches and facilitates accurate and efficient sequence comparison.

## Supplementary Note F: Tables

**Fig. E.3.**
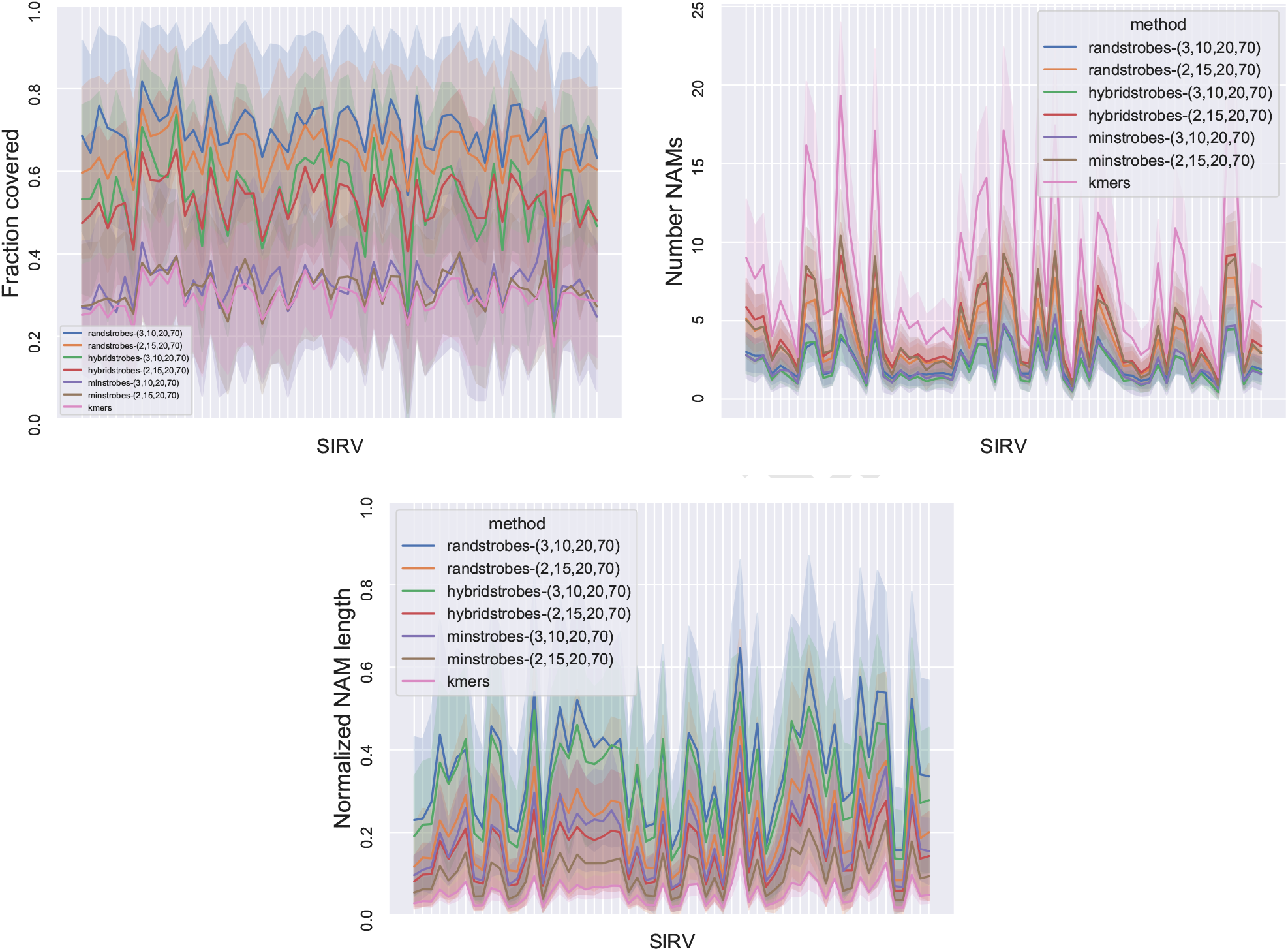
Comparison between strobemers and k-mers when matching 100 ONT cDNA reads to themselves from each of the 61 SIRVs. For each reference SIRV, there are 9,900 sequence mappings as read self-mapping results, which produce perfect matches are excluded. Each tick on the x-axis corresponds to a SIRV. Panel **A** shows total fraction of reference reads covered by NAMs from query reads (y-axis). Panel **B** shows the number of NAMs (y-axis) between the query and reference reads reads. Panel **C** shows the average normalized NAM length from all the NAM matches. The normalized NAM length is the length of the NAM divided by the length of the read acting as the reference in the given match. The line shows the mean and the shaded area displays the standard deviation of the reads. A high NAM coverage and low number of NAMs means long contiguous matches and facilitates accurate and efficient sequence comparison.

**Fig. E.4.**
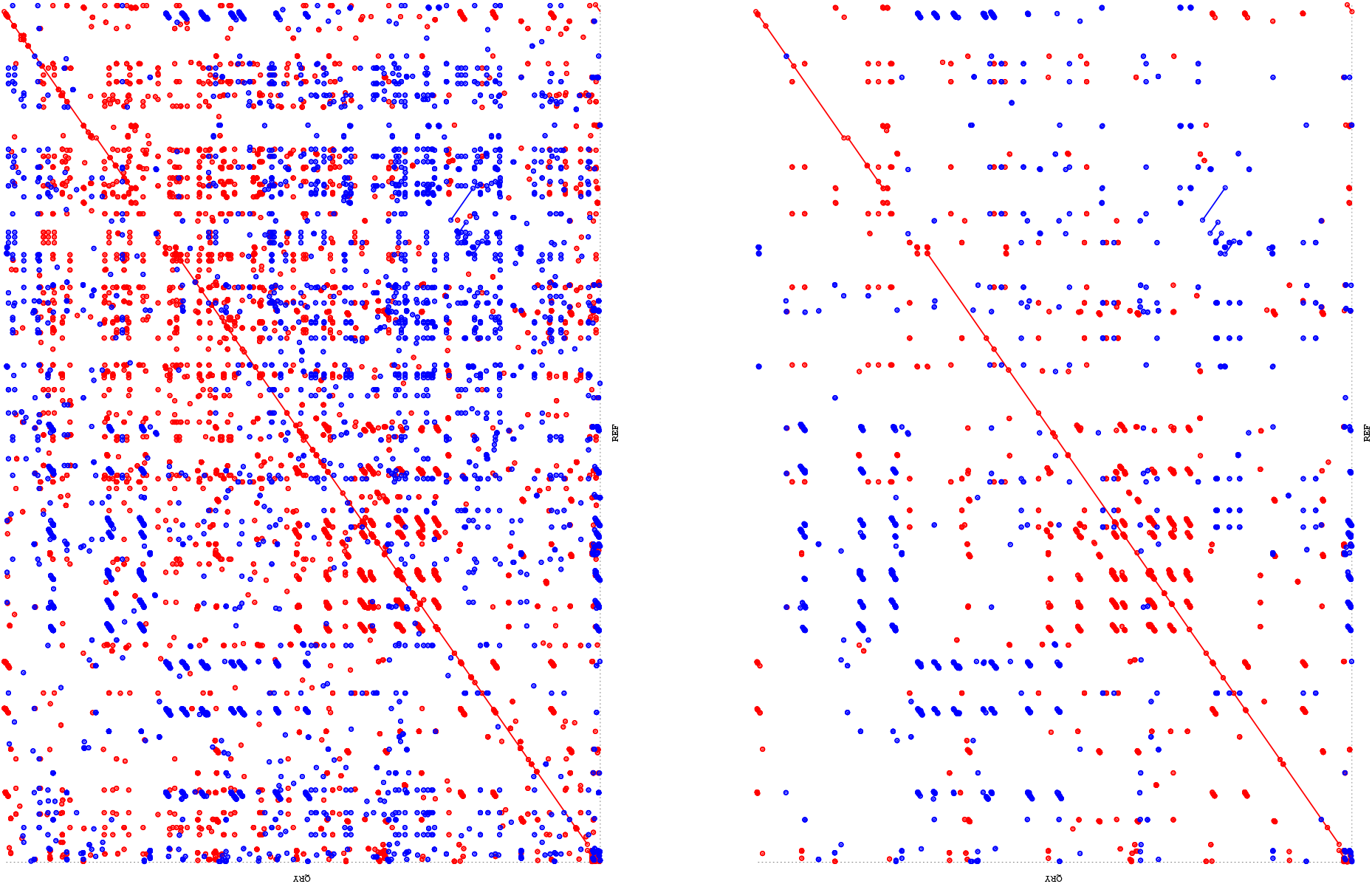
Dotplots of mapping two different *E. coli* genomes to each other using (**A**) hybridstrobes parametrized by (3,15,1,70), and (**B**) hybridstrobes parametrized by (2,15,20,120) with minimizer thinning protocol using *w* = 20.

